# Sniffing can be initiated by dopamine’s actions on ventral striatum neurons

**DOI:** 10.1101/2024.02.19.581052

**Authors:** Natalie L. Johnson, Anamaria Cotelo-Larrea, Lucas A. Stetzik, Umit M. Akkaya, Zihao Zhang, Marie A. Gadziola, Adrienn G. Varga, Minghong Ma, Daniel W. Wesson

## Abstract

Sniffing is a motivated behavior displayed by nearly all terrestrial vertebrates. While sniffing is associated with acquiring and processing odors, sniffing is also intertwined with affective and motivated states. The neuromodulatory systems which influence the display of sniffing are unclear. Here, we report that dopamine release into the ventral striatum is coupled with bouts of sniffing and that stimulation of dopaminergic terminals in these regions drives increases in respiratory rate to initiate sniffing whereas inhibition of these terminals reduces respiratory rate. Both the firing of individual neurons and the activity of post-synaptic D1 and D2 receptor-expressing neurons in the ventral striatum are also coupled with sniffing and local antagonism of D1 and D2 receptors squelches sniffing. Together, these results support a model whereby sniffing can be initiated by dopamine’s actions upon ventral striatum neurons. The nature of sniffing being integral to both olfaction and motivated behaviors implicates this circuit in a wide array of functions.

## Introduction

Sniffing is a volitional behavior displayed by nearly all terrestrial vertebrates and is defined by the rapid inhalation and exhalation of air through the nose. The commonplace function ascribed to sniffing is that it aids in perceiving odors^1,2^. This is supported by studies reporting that rodents increase their respiratory frequency from ∼2-4Hz at rest, up into an around 6-12Hz range when sampling an odor^3–6^. In addition to supporting olfaction, sniffing is also intertwined with affective and motivated states wherein sniffing is considered an appetitive behavior^7–9^. For instance, rodents, dogs, and even semi-aquatic vertebrates increase their sniffing frequency while foraging for food and in anticipation of reinforcers in instrumental tasks^5,6,9,10^. The frequency of sniffing also appears to influence emotional states^11,12^. Furthermore, sniffing serves as a “master clock” for essential physiological rhythms including whisking, wherein the rhythm of sniffing is coupled with whisking and also other head movements^13,14^. Therefore, it is clear that sniffing holds a ubiquitous position in nature.

The automatic coordination of respiration, and the tight control of the inspiratory / post-inspiratory / expiratory rhythm, are afforded by brainstem structures which elegantly time breaths, control their depth, and send these instructive signals toward the breathing plant by means of the spinal and cranial nerves^15,16^. While likely many of the respiratory brainstem structures that participate in the control of basal breathing are integral for the orchestration of sniffing, the exact mechanisms and neural basis for sniffing still remain unknown. Although sniffing is associated with increased bursting in the inspiratory rhythm generator, the pre-Bӧtzinger complex^17^, optogenetic stimulation of this region has yet to reproduce the rapid breathing rates that characterize sniffing^18–20^, unlike a subset of neurons in the pontine respiratory modulator, parabrachial nucleus^21^. Additionally, subsets of neurons in the expiratory control center, the retrotrapezoid nucleus, increase their activity shortly before the onset of sniffing^22^. These neurons provide input to the pre-Bӧtzinger complex^23^, as well as the facial motor nucleus, another region recognized to play a direct role in controlling sniffing^24^. While these findings together begin to uncover the circuitry for sniffing, more work needs to be done to pinpoint all the components, and specifically the neuromodulatory circuitry, which invigorates an animal to go from basal breathing which serves the purpose of gas exchange, to engage in the voluntary act of sniffing.

We reasoned the mesolimbic dopamine (DA) system as a prime candidate for the invigoration of sniffing. Electrical stimulation of the ventral tegmental area (VTA) elicits vigorous sniffing^25^. Neurons synthesizing DA in the VTA project into the basal forebrain, including into the ventral striatum. The ventral striatum, considered a limbic-motor interface^26^, is well-known to integrate cortical and limbic circuitry to adaptively guide behavior^27,28^. The major subcomponents of the ventral striatum, including the nucleus accumbens (NAc) and the tubular striatum (TuS; also known as the olfactory tubercle) are recipient of dense DAergic inputs from the VTA. Neurons in both the NAc and TuS are largely spiny projection neurons expressing D1- or D2-like receptors which mediate actions of DA^29–31^.

Through studies of DA’s role in the ventral striatum, we have come to learn that DA’s actions in the ventral striatum influence a wide-range of behaviors and states, such as aversion behaviors, and also cellular plasticity needed for learning^32–37^. DA in the ventral striatum is also a major mediator of motivated states and reinforcement with manipulation of DA, and D1 and D2 neural activity in the ventral striatum guiding those functions^36,38–40^. Since sniffing is a motivated behavior, it is reasonable to believe the ventral striatum’s role in motivational control is in-part a regulatory system which influences the occurrence of sniffing. Indeed, DAergic manipulations in the ventral striatum also influence exploratory behaviors in open-field and other gross motor behavior assays and recently it was reported that DAergic inputs regulate whisking^41^. Mesolimbic DA thus seems a quintessential neural system for supporting sniffing. Here, we investigated the role of mesolimbic DA in the display of sniffing behavior to find that sniffing can be initiated by DA’s actions upon ventral striatum neurons.

## Results

### Sniffing frequency is associated with mesolimbic DA release

We initially sought to understand the relationship between sniffing and DA release in the ventral striatum. To address this, we first performed a tracing study to identify ventral striatum subregions wherein DA inputs terminate for subsequent monitoring of DA during behavior. It is well known in both mice and rats that the ventral striatum receives dense input from mesolimbic DAergic neurons^28,42,43^ and that DA dynamics may differ between ventral striatum subregions^44,45^. In contrast to the innervation of the dorsal striatum and NAc, quantitative analyses of DA input to the TuS in mice is especially lacking. To address this, we injected AAV1-hSyn-FLEX-mGFP-2A-Synaptophysin-mRuby into the VTA of *DAT^IRES^-Cre* mice (**Fig 1A & Supplementary Fig 1**). This approach allowed for visualization of fluorescent puncta, which indicate DAergic presynaptic terminals, throughout the TuS and NAc (**Fig 1B**). The injections resulted in GFP expression throughout the VTA (**Supplementary Fig 1**). We quantified fluorescent puncta in the NAc Core (NAcC), NAc Shell (NAcSh), medial TuS (mTuS), and lateral TuS given different roles for these subregions in the regulation of motivated behaviors (**Fig 1C**)^46–49^. As expected, we observed robust fluorescence, indicating synaptophysin+ DAT VTA terminals, in the TuS and NAc, most notably in the anterior aspect of the mTuS and NAc (**Fig 1C**) with 113% and 78% more innervation in the anterior versus posterior halves of the mTuS and NAcSh, respectively (the anterior or posterior-most 200µm were used for quantification of these regions). Following this we focused analyses on the anterior aspect of both structures. While the mTuS was rich with synaptophysin+ DAT VTA terminals, this contrasted to the lateral TuS which was largely void of innervation (**Fig 1C & 1D**, rmANOVA *F*(1.52, 7.6)=31.74 *p*=0.0003, Tukey’s multiple comparisons ant mTuS vs. ant LatTuS *p*=0.014). Comparable amounts of synaptophysin+ DAT VTA terminals were found in the anterior NAcSh and anterior NAcC (**Fig 1C & 1D**, Tukey’s multiple comparisons antNAcSh vs. antNAcC, *p*=0.073). The density of input across the anteromedial aspects of the TuS, NAcSh, and NAcC were all similar (**Fig 1C & 1D;** Tukey’s multiple comparisons anterior mTuS vs. anterior NAcSh *p*=0.217; anterior mTuS vs. anterior NAcC *p*=0.878). Together this work adds to literature indicating that the anterior NAcSh, NAcC, and mTuS are the major recipients of VTA DAergic input in the ventral striatum and provide quantitative support for focusing on the medial aspects of these regions in mice.

**Figure 1.**
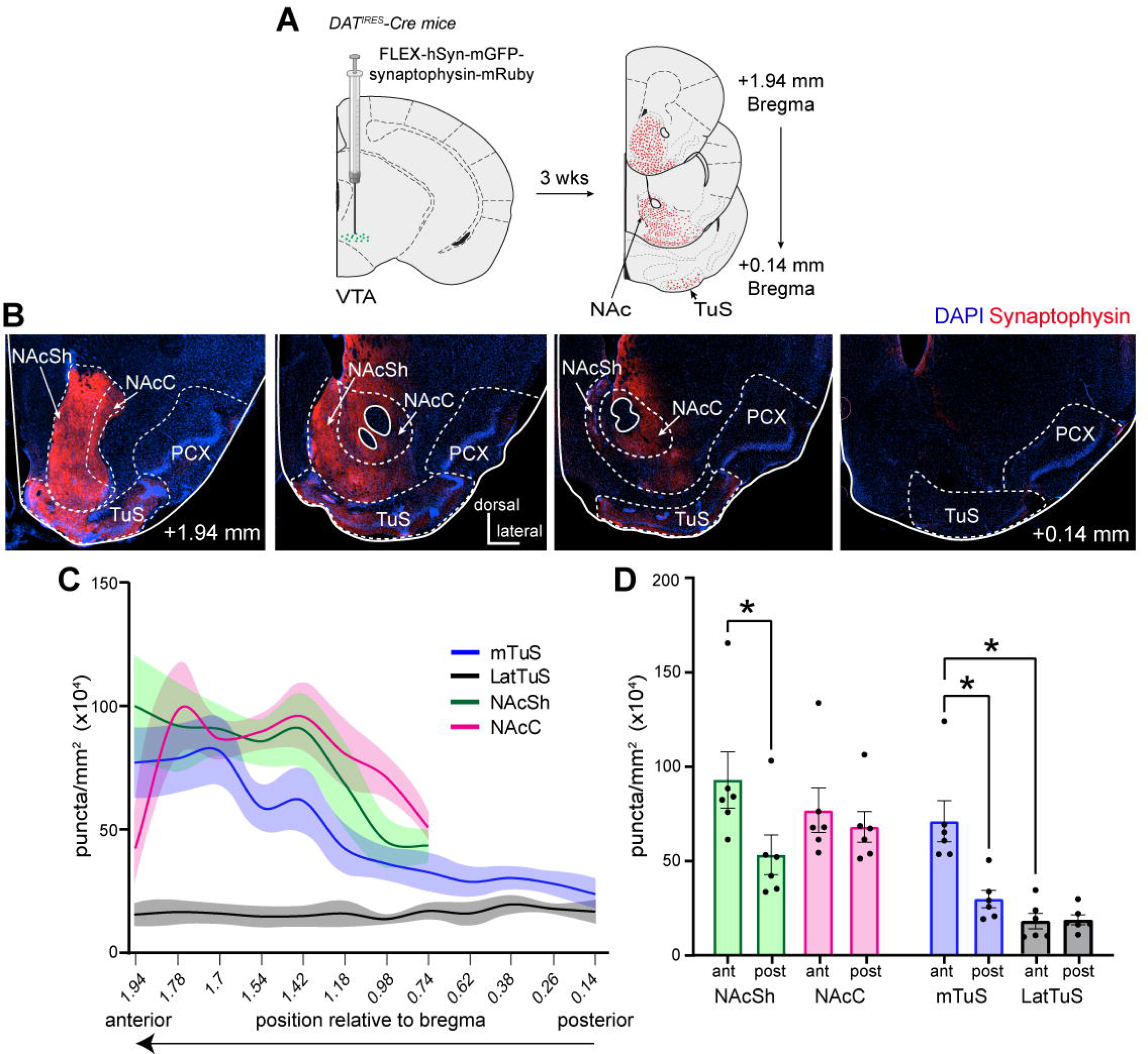
Organization of midbrain dopamine input into the tubular striatum and nucleus accumbens. **A.** Schematic of tracing paradigm. *DAT^IRES^ -Cre* mice were injected into their VTA with AAV.FLEX.hSyn.mGFP.synaptophysin.mRuby allowing for mGFP expression in DAT+ cell bodies and synaptophysin.mRuby+ puncta in the terminals of DAT+ cells. Approximately 3 weeks later, coronal sections throughout the ventral striatum were collected for later fluorescence imaging and quantification of synaptophysin.mRuby+ puncta. **B.** Example of synaptophysin.mRuby expression throughout the ventral striatum, with key regions of interest outlined by dashed lines. **C.** Quantification of synaptophysin+ puncta throughout the anterior-posterior spans of regions of interest (*n*=6 mice). Data [puncta #’s/mm^2^] are represented as mean ± SEM. Data smoothed for visualization purposes only. **D.** Histograms representing data in (**C**) with regions of interest segregated into their most anterior and posterior aspects (anterior most half vs. posterior most half). Data are mean ± SEM, points = individual mice. \**p*<0.5. abbreviations: VTA (ventral tegmental area), NAcSh (nucleus accumbens shell), NAcC (nucleus accumbens core), mTuS (medial aspect of the tubular striatum), LatTuS (lateral aspect of the tubular striatum), PCX (piriform cortex). (the anterior or posterior-most 200µm were used for quantification of these regions).

Having established this, we next sought to investigate the temporal relationship between mesolimbic DA release and sniffing behavior. We took advantage of whole-body plethysmography to assess respiratory cycles in awake mice and identify bouts of sniffing. In this method, breaths are detected by a pressure transducer which accesses the inside of the chamber (**Fig 2A**). The technical needs of our experiments require a patch cable attached to the mouse in the chamber (*e.g.,* **Fig 2C**) which may impair fidelity of the respiratory signal since the cable would need to exit the plethysmography chamber which is normally air-tight. Further, we wished to identify temporal relationships between sniffing and other signals (DA, photostimulation, *etc*) and this mandates that first we establish a temporal relationship between breaths detected with whole-body plethysmography and breathing of mice which occurs only through their noses since they are obligate nasal air breathers. Therefore, we simultaneously monitored respiration via the plethysmograph as well as by means of a chronically-implanted cannula into the dorsal nasal recess of awake mice (**Fig 2A**)^6^. For the intranasal measures, a flexible tube was connected to the nasal cannula which passed through the top of the plethysmograph via a snug 3D-printed fitting which allowed for mice to explore and freely-rotate within the plethysmograph. The end of this tube terminated into a pressure transducer for amplification of intranasal pressure changes. Using this simultaneous approach, we confirmed in three mice that whole-body plethysmography, even with the patch cable freely passing through the chamber, reliably detected intranasal respiratory cycles, and faithfully followed during high frequency sniffing bouts (**Fig 2Bi & 2Biii**). The only notable difference we observed for the purpose of this study was a slight temporal lag with the peaks of the whole-body plethysmograph signal trailing those of the intranasal signal (**Fig 2Bii**). This lag is anticipated since the origin of the signal is from within the mouse (intranasal) versus from within a chamber (plethysmograph). Across the three mice as they spontaneously transition in and out of sniffing bouts, the mean lag ranged from 0.028 ± 0.007s to .036 ± 0.0004s (using data with respiratory rates ranging from ∼2-12Hz, as in **Fig 3Biii**). While solely at resting respiration (2-4Hz), the lag between the two signals increased and ranged from 0.053 ± 0.001s to 0.10 ± 0.001s (across-animal range). Nevertheless, in events wherein respiratory cycles were detected in the nose, in >99% of the cases, they were also detected by the plethysmograph (3,239 breaths across three mice, a restricted range of data from one mouse shown in **Fig 2Bii**).

**Figure 2.**
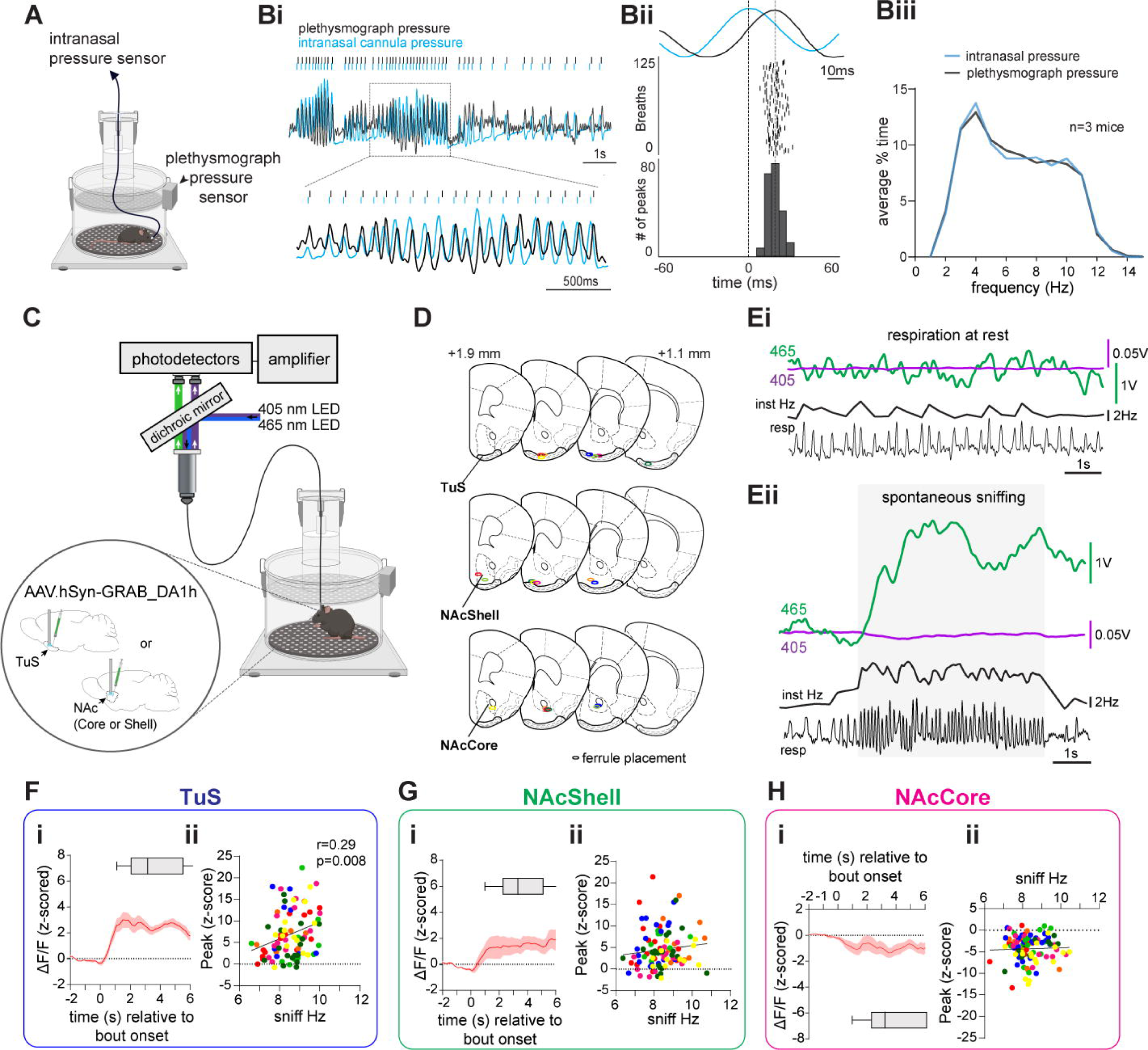
Self-motivated spontaneous sniffing is associated with DA levels. **A.** Diagram of the setup for simultaneous measurement of respiration by means of both whole-body plethysmography and an intranasal pressure cannula. Three mice were implanted with a cannula in its dorsal nasal recess and placed in the plethysmograph. Intranasal pressure changes were monitored by a pressure sensor connected to the cannula by a flexible tube passed through a fitted plug at the top of the plethysmograph and compared to whole-body changes. Image made with BioRender. **Bi**. Overlaid representative respiratory traces acquired from the plethysmograph (black) and intranasal cannula (blue) from an awake mouse. An example of high frequency sniffing is shown zoomed in and the expanded inset shows fidelity between the whole-body pressure and intranasal pressure signal. **Bii.** Example of the latency difference between respiratory peaks recorded by intranasal pressure and plethysmograph pressure during a high frequency sniffing bout with detected respiratory events (*n*=125 respiratory cycles). Overlaid respiratory traces (top) shows an intranasal pressure peaks aligned to time 0, and a plethysmograph pressure peak detected <60ms later. **Biii**. Histograms comparing respiratory frequencies measured from the plethysmograph and intranasal cannula. The plethysmograph reliably detected sniffing at both low and high respiratory frequencies with >99% accuracy (out of 3,239 respiratory events collected from 3 mice). **C.** Schematic of fiber photometry and plethysmograph system used to simultaneously record GRAB_DA_ signals and sniffing behavior in unrestrained mice. Animals were injected with AAV.hSyn-GRAB_DA1h and implanted with an optic fiber in the TuS, NAcSh, or NAcC. 3-5 weeks later, animals were placed in the plethysmograph and 465nm and 405nm light were passed through dichroic mirrors before reaching the animal via patch cable and implanted optical fiber. GRAB_DA_ and UV emissions were amplified by photoreceivers before being digitized along with simultaneous respiratory data. Image made with BioRender. **D.** Optic fiber implant locations in the TuS, NAcSh, and NAcC (*n*=7 mice/group). Within each region, ellipse color corresponds to same colored data in **Figures 2F-H and 3C-H**. **E.** Example of raw 465nm GRAB_DA_ (green) and 405nm UV (purple) signals recorded in the TuS during a spontaneous sniff bout. Raw respiratory trace (resp) is shown below with interpolated instantaneous frequency above (Inst Hz-black trace). Sniff duration is indicated by the grey shaded box. **Fi.** Averaged z-scored ΔF/F traces show an increase in GRAB_DA_ levels in TuS during spontaneous sniff bouts. Average bout duration indicated by overlaid box plot. **Fii**. Significant positive correlation between GRAB_DA_ levels in the TuS (the peak z-score during a spontaneous sniff bout) and average sniff frequency during bout (*r*=0.29, *p*=0.008). **Gi**. Averaged z-scored ΔF/F traces similarly show an increase in GRAB_DA_ levels in the NAcSh during spontaneous sniff bouts. Average bout duration indicated by overlaid box plot. **Gii**. There was no correlation between GRAB_DA_ levels in the NAcSh during spontaneous bout and sniffing frequency (*r*=0.12, *p*=0.249). **Hi**. Conversely, averaged z-scored ΔF/F traces show a decrease in GRAB_DA_ levels in the NAcC during spontaneous sniff bouts. Average bout duration indicated by overlaid box plot. **Hii**. There was no significant correlation between GRAB_DA_ levels in the NAcC during spontaneous bout and sniffing frequency (*r*=0.038, *p*=0.703). Abbreviations: resp (respiratory trace), Inst Hz (instantaneous frequency). Data in (Fi), (Gi), and (Hi) smoothed for visualization purposes only.

**Figure 3.**
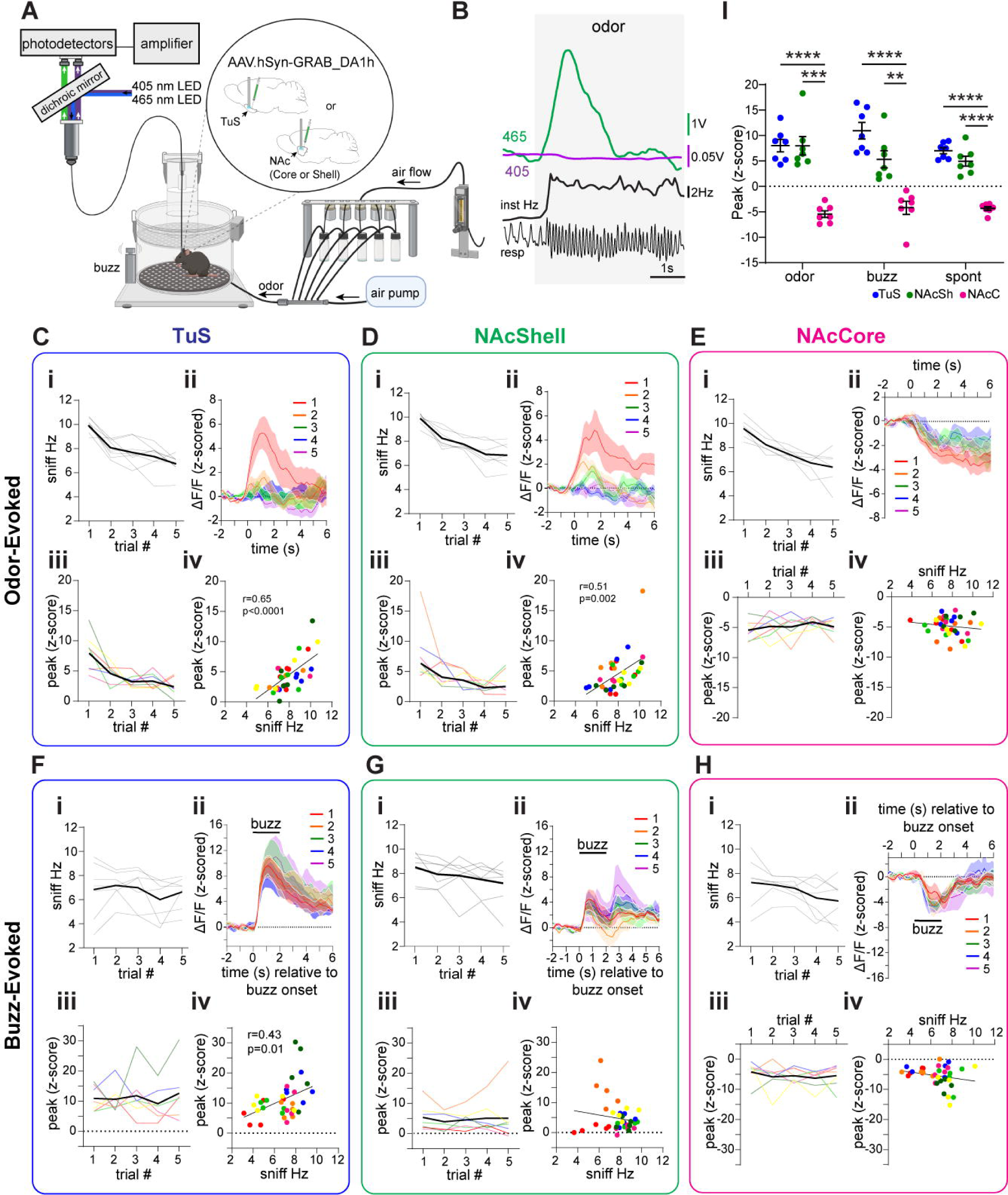
Bidirectional dopamine dynamics upon sensory-driven sniffing. **A.** Schematic of fiber photometry and plethysmograph system used to simultaneously record GRAB_DA_ signals in the TuS, NAcSh, and NAcC and sniffing behavior in unrestrained mice (*n*=7/group) as used in Fig 2. Additionally, an olfactometer was used to deliver odors to animals in the plethysmograph, and a vibration motor (buzz) was attached to the outside of the plethysmograph. Image made with BioRender. **B.** Example of raw 465nm GRAB_DA_ (green) and 405nm UV (purple) signals recorded in the TuS during an odor-evoked bout. Raw respiratory trace (resp) is shown below with interpolated instantaneous frequency above (Inst Hz-black trace). Grey shaded box denotes odor analysis window, starting 3 seconds following odor onset to account for the delay of odor delivery to the plethysmograph. **C-E.** Odor-evoked sniffing behavior and GRAB_DA_ signals from mice with injections and optic ferrule implants in the TuS (C), NACSh (D), and NAcC (E). Average instantaneous sniffing frequency of TuS (Ci), NAcSh (Di), and NAcC (Ei) implanted mice across repeated odor presentations. Mean shown in black with individual mouse data across trials shown in light grey. Averaged z-scored ΔF/F across all 5 odor presentations in the TuS (Cii), NAcSh (Dii), and NAcC (Eii). Averaged peak of the z-scored GRAB_DA_ response across trials in the TuS (Ciii), NAcSh (Diii), and NAcC (Eiii). Mean shown in black with individual mouse data across trials color coded to ellipse colors shown in **Figure 2D**. Significant positive correlations between GRAB_DA_ levels (peak z-score during odor-evoked activity) and average sniff frequency during bout in the TuS, (*r*=0.65, p<0.0001, Civ) and NAcSh (*r*=0.51, p=0.002, Div) but not in the NAcC (*r*=-0.14, *p*=0.414, Eiv). **F-H.** Buzz-evoked sniffing behavior and GRAB_DA_ signals from same mice with injections and implants in the TuS (F), NAcSh (G), and NAcC (H). Average instantaneous sniffing frequency of TuS (Fi), NAcSh (Gi), and NAcC (Hi) implanted mice across repeated odor presentations. Mean shown in black with individual mouse data across trials shown in light grey. Averaged z-scored ΔF/F across all 5 odor presentations in the TuS (Fii), NAcSh (Gii), and NAcC (Hii). Horizontal line depicts 2s buzz duration. Averaged peak of the z-scored GRAB_DA_ response across trials in the TuS (Fiii), NAcSh (Giii), and NAcC (Hiii). Mean shown in black with individual mouse data across trials color coded to ellipse colors shown in **Figure 2D**. Significant positive correlation between GRAB_DA_ levels and sniffing frequency in the TuS, (*r*=0.43, *p*=0.011, Fiv), but not the NAcSh (during buzz: *r*=-0.17, *p*=0.316; upon buzz offset: *r*=-0.074, *p*=0.673, Giv) or NAcC (*r*=-0.22, *p*=0.194, Hiv). **I.** Comparison of the peak-evoked GRAB_DA_ responses across the TuS, NAcSh, and NAcC during odor-, buzz-, and spontaneously-evoked sniffing. Data are represented as mean ± SEM. Points = individual mice. \*\*\*\**p*<0.0001, \*\*\**p*<0.001, \*\**p*<0.01. Data shown in (Cii), (Dii), (Eii), (Fii), (Gii), and (Hii) are smoothed for visualization purposes only.

Having confirmed the ability to accurately detect both individual respiratory cycles and sniffing bouts alike, we next monitored the relationship between DA levels in the ventral striatum as they relate to sniffing. We injected mice into either the medial TuS, the NAcSh, or NAcC with an AAV encoding GRAB_DA_ (pAAV.hSyn-GRAB_DA1h)^50^ and in the same surgery implanted into the structure receiving the AAV, a 400µm optical fiber for later fiber photometry. Other than for the purpose of showing example traces, for other analyses we subtracted the isosbestic 405nm signal from the GRAB_DA_ signal. Following several weeks for surgical recovery and viral expression, we placed the mice in a plethysmograph for simultaneous monitoring of respiration and DA levels (**Fig 2C**). Since mice rapidly transition in and out of sniffing bouts^6^, we restricted our analyses to more stable/pronounced bouts versus short bouts by only defining a sniff bout as the occurrence of sniffing ≥6Hz for ≥1s in duration. Further, in some instances, mice continuously sniff for numerous seconds and then reengage in another bout shortly thereafter. Since these may reflect periods of more generalized arousal versus spontaneous exploratory behavior, we excluded sniff bouts occurring in proximity to these long (>10s) sniffing bouts. We analyzed the relationship between DA levels and sniffing in several manners, and in all analyses, we accounted for the lag between when a nasal inhalation occurs and when the plethysmograph detects the inhalation by subtracting the conservative 28ms lag (see **Fig 2Bii**) from the onset event of every peak-detected respiratory cycle.

We analyzed DA levels across trials wherein mice in the plethysmograph spontaneously sniffed, were delivered odors to evoke sniffing, or were delivered an auditory/somatosensory stimulus by connecting a vibration motor to the plethysmograph which modestly resonated the chamber while simultaneously providing a small auditory ‘buzz’ cue due to its motor (**Supplementary Fig 2**). All mice were confirmed to have implants extending into the mTuS, NAcC, or the NAcSh (**Fig 2D**).

First, we looked at the relationship between spontaneous sniffing like that displayed by mice during self-motivated states of exploration. As shown in the example traces from a mouse implanted into the TuS (**Fig 2E**), whereas no detectable changes in DA levels were observed when mice were simply breathing at baseline frequencies (‘respiration at rest’; **Fig 2Ei & Supplementary Fig 3A**), spontaneous sniffing was associated with elevations in the GRAB_DA_ signal (**Fig 2Eii**). Across mice, GRAB_DA_ levels during spontaneous sniffing were elevated in both the TuS and the NAcSh, but in contrast were decreased in the NAcC (**Fig 2Fi, 2Gi, 2Hi**; TuS paired *t*(6)=11.85 *p*<0.0001; NAcSh paired *t*(6)=6.56, *p*=0.0006; NAcC paired *t*(6)=12.15, *p*<0.0001). Only in the TuS, was the average frequency of the spontaneous sniffing bout correlated with the amplitude of the GRAB_DA_ signal (**Fig 2Fii, 2Gii, & 2Hii**; TuS Pearson’s *r*(83)=0.29, *p*=0.008), but not in the NAcSh or NAcC (NAcSh Pearson’s *r*(101)=0.12, *p*=0.2493; NAcC Pearson’s *r*(102)=0.038, *p*=0.703). Notably, neither the isosbestic 405nm signal acquired from these same animals, nor the DA-insensitive control virus, GRAB_DA_-mut, acquired in a separate cohort of 9 mice, showed comparable changes during spontaneous sniffing (**Supplementary Fig 4**).

Next, we investigated the relationship between odor-evoked sniffing and DA levels (**Fig 3A**). Each mouse was presented with pseudorandom trials of odors, including R(+)-limonene, thioglycolic acid, and peanut oil, each five times across a behavior session. These odors were selected based upon their known positive hedonic qualities perceived by mice, in order to elicit vigorous sniffing ^51–53^. As shown in the example in **Figure 3B** from a mouse implanted in the TuS, the first presentation of thioglycolic acid was associated with robust increases in sniffing, which was accompanied by an increase in the GRAB_DA_ signal. Across all groups (TuS, NAcSh, NAcC), sniffing frequency was elevated compared to baseline for the first presentation of odors (TuS paired *t*(6)=19.23, *p*<0.0001; NAcSh paired *t*(6)=13.27, *p*<0.0001; NAcC paired *t*(6)=14.35, *p*<0.0001), which habituated across subsequent presentations (TuS rmANOVA *F*(2.26,13.55)=14.64, *p*=0.0003; NAcSh rmANOVA *F*(2.44, 14.65)=32.65, *p*<0.0001; NAcC rmANOVA *F*(1.59, 9.55)=23.5, *p*=0.0003) (**Figs 3Ci, 3Di, & 3Ei**). Simultaneously during novel odor-evoked sniffing bouts (trial #1 of odor), we observed elevations in the GRAB_DA_ signal in both the TuS and NAcSh (**Fig 3Cii & 3Dii**; TuS paired *t*(6)=6.54, *p*=0.0006; NAcSh paired *t*(6)=4.38, *p*=0.005). The increases rose soon around sniff bout onset and decayed within a few seconds. In contrast, the NAcC displayed reduction in the GRAB_DA_ signal upon sniffing (**Fig 3Eii**; paired *t*(6)=8.35, *p*=0.0002).

The elevation in the GRAB_DA_ signal was not solely due to odor input since the signal became attenuated across repeated odor deliveries in the TuS and NAcSh **(Fig 3Ciii & 3Diii**; trials #1-5; TuS rmANOVA *F*(2.08, 12.48)=7.942, *p*=0.006; NAcSh rmANOVA: *F*(1.90,11.39)=5.563, *p*=0.022), whereupon trial 5, the GRAB_DA_ signal was observed near baseline levels. In the NAcC, reductions in the GRAB_DA_ signal was not relieved with repeated odor presentations (**Fig 3Eiii**; rmANOVA *F*(1.98, 11.90)=0.5760, *p*=0.58). Also supporting that the change in amplitude of the GRAB_DA_ signal is related to sniffing, we found that the frequency of odor-evoked sniffing correlated with the peak of the GRAB_DA_ signal in both the TuS and NAcSh (TuS Pearson’s *r*(34)=0.65, *p*<0.0001; NAcSh Pearson’s *r*(34)=0.51, *p*=0.002), but not in the NAcC (Pearson’s *r*(34)=-0.14, *p*=0.414) (**Figs 3Civ, 3Div, & 3Eiv**). Interestingly here, the odor-evoked sniffing in the NAcSh was correlated with GRAB_DA_ levels which was not the case with spontaneous sniffing (**Fig 2G**), suggesting in the NAcSh the relationship may be dependent upon odor.

Finally, we assessed the impact of the multimodal buzz stimulus on GRAB_DA_ levels. While not as dramatic nor reliable as an odor (as in **Fig 3Ci, 3Di, & 3Ei**), the buzz stimulus evoked sniffing greater than baseline in the majority of mice (**Fig 3Fi, 3Gi, & 3Hi**). Correspondingly, DA was elevated upon initial buzz-evoked sniffing in the TuS and the NAcSh (**Figs 3Fii & Gii**; TuS paired *t*(6)=6.7*7, p*=0.0005; NAcSh paired *t*(6)=3.17, *p*=0.019). The buzz-evoked sniffing did not strongly habituate across repeated trials, and likewise, GRAB_DA_ levels also remained consistent across trials (**Fig 3Fi & 3Fiii**, **Fig 3Gi & 3Giii**; TuS sniffing rmANOVA *F*(2.57, 15.4_)_=1.351, *p*=0.293; TuS GRAB_DA_ rmANOVA *F*(1.90, 11.38)=0.49, *p*=0.616; NAcSh sniffing rmANOVA *F*(2.43, 14.58)=2.97, *p*=0.0752; NAcSh GRAB_DA_ rmANOVA *F*(1.32, 7.94)=0.27, *p=*0.6786). Interestingly, in the NAcSh, there was a biphasic increase in GRAB_DA_ with the first phase occurring upon buzz onset, and the second upon buzz offset (**Fig 3Gii**). Whereas in the NAcSh, GRAB_DA_ levels measured upon either phase were not correlated with the frequency of sniffing (**Fig 3Giv**; during buzz Pearson’s *r*(34)=-0.17, *p*=0.316; upon buzz offset Pearson’s *r*(34)=-0.074, *p*=0.673), the GRAB_DA_ levels upon buzz strongly correlated with frequency of sniffing in the TuS (**Fig 3Eiv**; Pearson’s *r*(34)=0.43, *p*=0.011). Similar to that seen for both spontaneous sniffing and odor-evoked sniffing, GRAB_DA_ levels in the NAcC were reduced upon buzz-evoked sniffing (**Fig 3Hii**; paired *t*(6)=3.23, *p*=0.018) which similarly did not habituate across trials (**Fig 3Hii & 3Hiii**; rmANOVA *F*(2.19, 13.14)=0.44, *p*=0.668). There was no correlation between buzz-evoked sniffing frequency and GRAB_DA_ levels in the NAcC (**Fig 3Hiv**; Pearson’s *r*(34)=-0.22, *p*=0.194).

When comparing the peak GRAB_DA_ responses across all regions and all conditions, we found similar evoked responses in the TuS and NAcSh to novel odor (trial #1 of odor) presentations and during spontaneous sniffing **(Fig 3I**; two-way rmANOVA, main effect of region *F*(2,18)=58.45, *p*<0.0001; Tukey’s multiple comparisons TuS vs. NAcSh odor *p*=0.99; TuS vs. NAcSh spont *p*=0.219). As with spontaneous sniffing, neither odor-nor buzz-evoked sniffing resulted in comparable changes in the separate cohort of 9 mice wherein we recorded GRAB_DA_-mut signal (**Supplementary Fig 4**). Taken together, these results indicate that sniffing often corresponds with increases in DA release in two specific sub-regions of the ventral striatum, and that the level of DA correlates to the frequency of sniffing.

### DA release into the ventral striatum elicits sniffing

Given the relationship between DA release in the ventral striatum and the occurrence and frequency of sniffing, we next sought to determine whether this release of DA is sufficient to elicit sniffing. To assess this, we injected a Cre-dependent Channelrhodopsin2 (AAV5-EF1α-DIO-hChR2(E123T/T159C)-EYFP) unilaterally into the VTA of *DAT^IRES^-Cre* mice (**Supplementary Fig 5**) and implanted a 300µm optical fiber terminating over the TuS, NAcSh, or NAcC to drive terminal release of DA upon light stimulation (**Fig 4A & 4B**). Control mice were injected with AAV solely expressing EYFP in the VTA (AAV5-EF1α-DIO-eYFP) and also received an optical fiber terminating over the TuS, NAcSh, or NAcC. One second long, 25Hz 473nm light stimulation was delivered through the optical fiber while mice were in the plethysmograph. This 25Hz stimulus was selected based upon prior work using a similar stimulus paradigm to elicit VTA DA release into the ventral striatum^39,40,54^. As shown in **Figure 4C**, in an example mouse, photostimulation of DAergic terminals in the TuS evoked a brief bout of sniffing. In this example mouse, the average latency to onset of a sniff bout was 240 ± 22.5ms (across trial range: 130-330ms; bout onset defined as the moment of achieving ≥6Hz respiration). Further, the average amount of time spent sniffing (time ≥6Hz) during photostimulation was 672 ± 59.6ms (across trial range: 310-1000ms). The light evoked sniffing in this example mouse was also reliable across repeated trials (**Fig 4C**).

**Figure 4.**
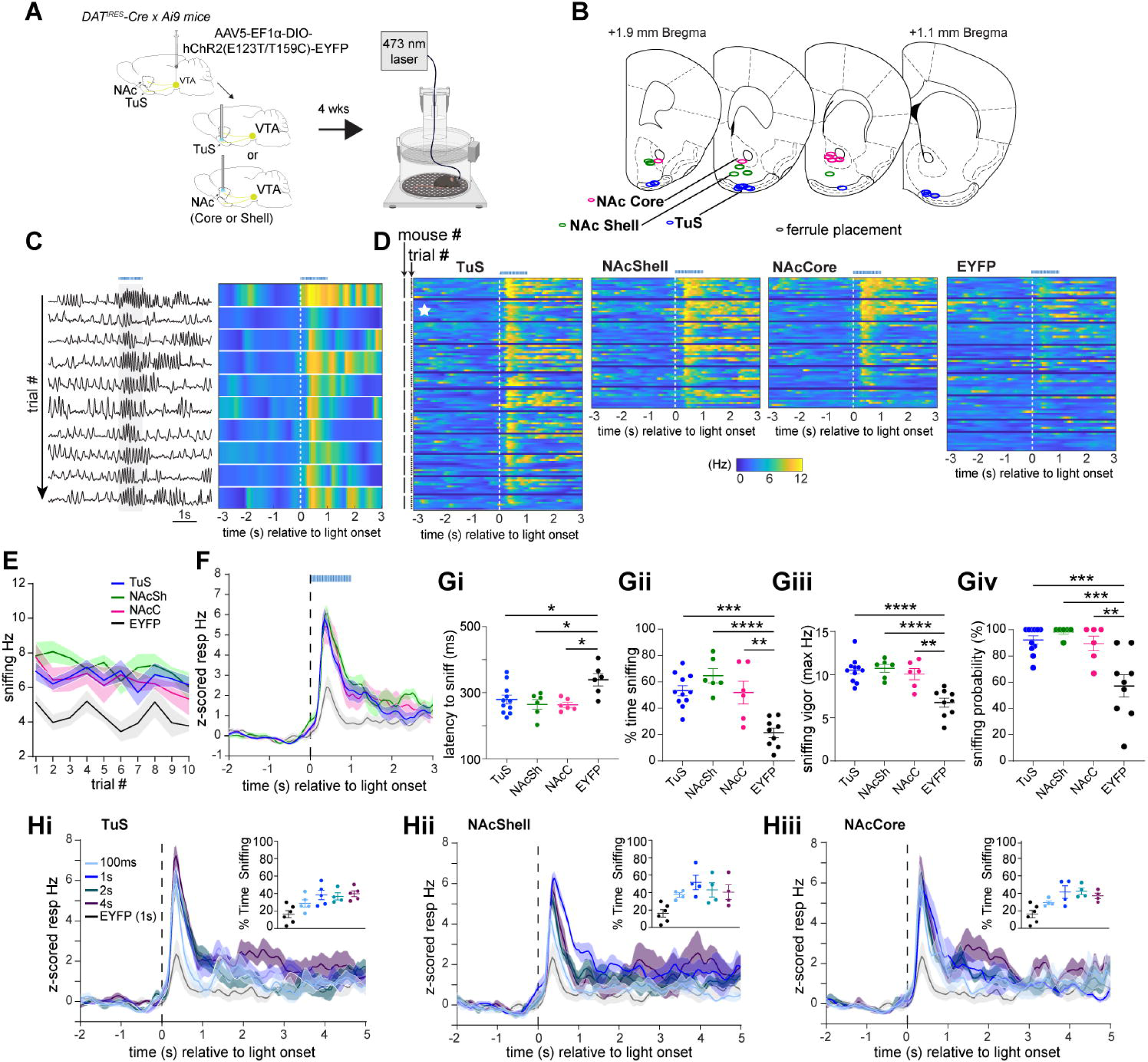
Initiation of sniffing by dopamine release into the ventral striatum. **A.** Schematic of optogenetically-evoked sniffing paradigm. *DAT^IRES^-Cre x Ai9* mice were injected with AAV-EF1α-DIO-hChR2(E123T/T159C)-EYFP in the VTA, and optic fibers terminating in the TuS, NAcSh or NAcC were implanted in the same surgery. Control mice were injected with AAV-Ef1α-DIO-EYFP. At least 4 weeks later, mice were tethered and placed in the plethsymograph to optogenetically stimulate DA terminal release while monitoring sniffing. Image (right) made with BioRender. **B.** Optic fiber implant locations in the TuS, NAcSh, and NAcC (*n*=6-11 mice/group). **C.** Example respiratory data from a TuS implanted mouse across 10 trials of light stimulation. Timing of the 1s long, 25Hz photostimulation depicted by grey shaded box and dotted blue horizontal line. Raw respiratory traces are shown on the left, and corresponding instantaneous frequency for each trial shown in 2-dimensional histograms on the right. Dotted white line in 2-dimenional histogram depicts stimulation onset. **D**. 2-dimensional histograms across all mice tested within each group, showing individual trial data within mice. Example data from (C) denoted by the white star. Separation between mice marked by dark blue line. Stimulation onset is denoted by the vertical dotted white line, and 1s, 25Hz stimulation shown by the dotted blue trace. **E.** Mean sniffing frequency ± SEM during photostimulation across 10 trials in each group. **F.** Averaged z-scored respiratory frequency ± SEM of photostimulation-evoked sniffing. Stimulation onset denoted by vertical dotted black line and 25Hz, 1s duration by horizontal blue line. Group color legend matches what is shown in (E). **G**. Parameters of optogenetically-evoked sniffing during 1s photostimulation. All data presented as mean ± SEM, points = individual mice. *\*\*\*\*p*<0.0001, \*\*\**p<*0.001, \*\**p<*0.01. **Gi.** Mean latency to sniff across groups. Animals that failed to reach 6Hz on >40% of trials were excluded from the EYFP control group. **Gii**. Mean percentage of time spent sniffing across groups. **Giii.** Max sniffing frequency reached during photostimulation. **Giv**. Mean probability that photostimulation will evoke sniffing. **H.** Averaged z-scored respiratory frequency during different photostimulation durations (100ms, 1s, 2s, 4s) compared to responses of EYFP controls stimulated for 1s in the TuS (Hi), NAcSh (Hii), and NAcC (Hiii). Insets in each panel show the percentage of time across all groups, calculated in a 4s window, that mice spend sniffing upon photostimulation.

In all three ChR2-injected groups of mice, we observed a rapid increase in respiratory frequency upon photostimulation (**Fig 4D**). This photostimulation-evoked display of sniffing was largely maintained across repeated trials (**Fig 4D & 4E;** testing for stability in Hz across trials: mixed-effects analysis *F*(4.09, 85.87)=2.34, *p*=0.058). While anticipated, EYFP control mice displayed a slight increase in respiratory frequency during photostimulation, likely due to the arousing nature of the visual stimulus leaking from the fiber implant. Alternatively, given that light stimuli can evoke DA release in the ventral striatum^55^ it is also possible that light-triggered DA release could have contributed to the increase in respiratory frequency seen in EYFP controls. Nonetheless, the frequency of the sniffing bout was lower in EYFP mice than in all ChR2-treated groups (**Fig 4D, right, Fig 4E & F**). As shown in **Figure 4F**, the temporal dynamics of photostimulation-evoked sniffing in ChR2-injected mice was remarkedly similar. The average latency to onset of a sniff bout was 279.4 ± 13.3ms SEM (inter-mouse range: 225-362.5ms) for TuS, 264.7 ± 14.83ms SEM (inter-mouse range: 202.5-299ms) for NAcSh, and 263.3 ± 7.52ms SEM (inter-mouse range: 242.2-290ms) for NAcC (**Fig 4Gi**). The average latency to the onset of a sniff bout in EYFP controls (339.4 ± 18.72ms SEM, inter-mouse range: 274.2-405ms) was significantly greater than in the ChR2-treated groups (all groups with means <290ms as in **Fig 4Gi**; one-way ANOVA *F*(3, 25)=5.11, *p*=0.007, Tukey’s multiple comparisons test TuS vs. EYFP *p*=0.0273, NAcSh vs. EYFP *p*=0.0141, NAcC vs. EYFP *p*=0.122). Further, the average amount of time spent sniffing during photostimulation was 534.1 ± 37.8ms (inter-mouse range 314.3-743ms) for TuS, 647.7 ± 53.7ms (inter-mouse range 479-834ms) for NAcSh, and 520.1 ± 85.7ms (inter-mouse range 254.4-755ms) for NAcC. All ChR2-treated mice spent more time sniffing than EYFP controls, and there were no significant differences between ChR2-treated groups (**Fig 4Gii**) (one-way ANOVA *F*(3,28)=14.02, *p*<0.0001; Tukey’s multiple comparisons test TuS vs. EYFP *p*=0.0001, NAcSh vs. EYFP *p*<0.0001, NAcC vs. EYFP *p*=0.0017). Further, the frequency of photostimulation-evoked sniffing (measured as the peak Hz during photostimulation) did not differ between ChR2-treated groups and was significantly greater than EYFP controls (**Fig 4Giii**; one-way ANOVA *F*(3, 28)=14.17, *p*<0.0001; Tukey’s multiple comparisons test TuS vs. EYFP *p*<0.0001, NAcSh vs. EYFP *p*<0.0001; NAcC vs. EYFP *p*=0.0010). Finally, we explored the reliability whereby a DAergic terminal photostimulation would elicit a sniffing bout. The probability that photostimulation elicited sniffing was robust, and significantly greater than in EYFP controls – indeed with many animals in each ChR2-treatment group displaying a sniff bout to *every* photostimulation trial and the vast majority to >80% of trials (**Fig 4Giv**; one-way ANOVA *F*(3, 28)=11.01, *p*<0.0001; Tukey’s multiple comparisons TuS vs. EYFP *p*=0.0002, NAcSh vs. EYFP *p*=0.0002, NAcC vs. EYFP *p*=0.004).

Since VTA neurons are known to co-release glutamate in addition to dopamine^35,56,57^, and knowing that ventral striatum neurons express glutamatergic receptors, we next sought to isolate the influence of the above effect (**Fig 4**) on DA versus glutamate. We crossed *VGluT2 fl/fl* mice with mice expressing Cre recombinase in all cells expressing tyrosine hydroxylase (*TH-Cre*), the rate-limiting enzyme upstream for DA synthesis. This approach ensures that *VGluT2* is lacking from tyrosine hydroxylase expressing neurons, thus eliminating the possibility of glutamate co-release from TH+ neurons. As above, we injected *VGluT2fl/fl* x *TH-Cre* mice with the same ChR2-expressing virus or associated control virus unilaterally into the VTA and implanted an optical fiber terminating over the TuS to drive terminal excitation. Since photostimulation in all three ventral striatum subregions of ChR2-expressing mice in the prior experiment drove sniffing, here we excluded the NAc groups for simplicity. As shown in **Supplementary Figure 6**, photostimulation of DAergic terminals in *VGluT2fl/fl* x *TH-Cre* mice elicited sniffing with comparable temporal dynamics, frequency, and probability as in *DAT^IRES^-Cre* mice (**see Fig 4**) and at greater amounts in *VGluT2fl/fl* x *TH-Cre* mice than in EYFP controls.

We next wanted to determine whether DA release in the ventral striatum serves to ‘initiate’ sniffing, or if it further can persistently drive sniffing over extended time courses. To test this, in the same ChR2-treated *DAT^IRES^-Cre* mice as in **Figures 4B-G**, we delivered repeated presentations of the 25Hz photostimulation for either 120ms, 1s, 2s, or 4s. We predicted that if DA initiates sniffing we would see transiently-evoked sniffing across all stimulation durations, whereas if DA drives sniffing, we would see the duration of sniffing bouts track the duration of the photostimulation. As shown in **Figure 4H**, we found that the dynamics of the sniffing bout were equivalent regardless of photostimulation duration. Indeed, in all three ChR2-treated groups, photostimulation of DAergic terminals elicited a statistically similar frequency of sniffing (TuS mixed-effects analysis; *F*(1.78, 6.53)=0.94 *p*=0.426; NAcSh rmANOVA *F*(1.63, 4.89)=0.924, *p*=0.436; NAcC rmANOVA *F*(1.93, 5.79)=0.24, *p*=0.789), and similar percentage of time sniffing (TuS mixed-effects analysis *F*(1.18, 4.32)=1.57, *p*=0.284; NAcSh rmANOVA *F*(1.50, 4.5)=0.77 *p*=0.479; NAcC rmANOVA *F*(1.43, 4.30)=1.81, *p*=0.404). These results indicate that DA release into the ventral striatum initiates sniffing, but that it does not persistently drive a sniff bout.

Stimulation of respiratory brain stem regions known to support breathing can generate sniffing patterns even in anesthetized mice (*e.g.* ^15,16,58^). Is mesolimbic DA input to the ventral striatum capable of initiating sniffing even in anesthetized mice? To test this, following the above experiments, *DAT^IRES^-Cre* mice injected with ChR2 were deeply anesthetized with urethane and photostimulation was delivered. The baseline respiratory pattern and rate in these mice were both stable and slow (<1.25Hz), indicating deep sedation. Breathing rate under anesthesia showed no change upon photostimulation of DAergic terminals in either the TuS or the NAcSh and NAcC (**Fig 5A - C**), suggesting that the VTA→ventral striatum DAergic circuit is a component of the voluntary, not automatic, respiration-control system.

**Figure 5.**
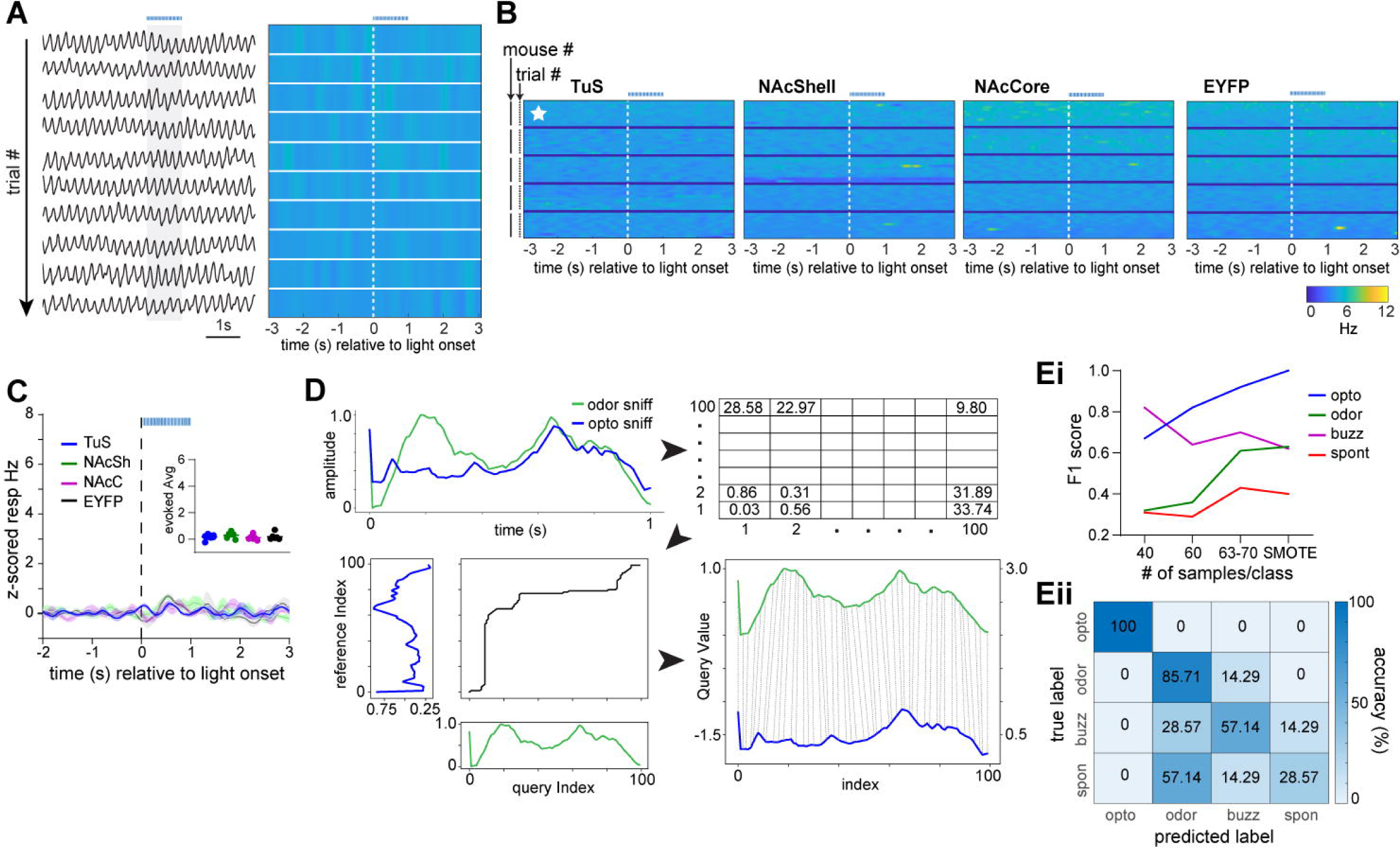
State-dependence of a unique dopamine-evoked sniffing pattern. **A.** Example respiratory behavior of a TuS implanted mouse under urethane anesthesia. Raw respiratory traces (left) and corresponding 2-dimensional histogram (right) across 10 trials of photostimulation. 1s long, 25Hz photostimulation duration depicted by grey shaded box and dotted blue horizontal line. Dotted white line in the 2-dimensional histogram = photostimulation onset **B.** 2-dimensional histograms across 5 representative mice per group showing individual trial data within mice. Starred data represents example data from (A). Separation between mice denoted by dark blue line. Stimulation onset is denoted by the vertical dotted white line, and 1s, 25Hz stimulation shown by the dotted blue trace. Color scale matches that of **Figures 4C and D**. Little to no change in coloring highlights the substantial lack of any modulation in breathing rate under anesthesia. **C.** Averaged z-scored respiratory frequency of photostimulation-evoked sniffing. Stimulation onset denoted by vertical dotted black line and duration by horizontal blue dotted line. Inset shows the evoked z-scored average across all groups during light stimulation. **D.** Overview of machine learning applied to respiratory traces starting with dynamic time warping (DTW) between a 1s odor-evoked sniff bout and 1s opto-evoked sniff bout. Bin size=10ms. 1s-long respiratory traces (100Hz) were converted to instantaneous frequency traces (also 1s long) and subjected to DTW to assess distance in proximal time-points. Subsequently, a model was trained on experimenter-identified traces from either buzz-, odor-, spontaneous-, or photostimulation-evoked respiration that were also previously subjected to DTW (see Methods). **Ei.** F1 score (single metric of model performance) for sniff types based on number of samples used per class. SMOTE=70 samples. **Eii.** Confusion matrix from SMOTE analyzed data displaying the classifier accuracy in predicting optogenetic-, odor-, buzz-, or spontaneously-evoked sniffing bouts. Classifier was trained on 80% of samples and tested on remaining 20%.

Sniffing bouts can be displayed in response to a variety of stimuli an animal may perceive in its environment. While not well-studied, it is possible the temporal dynamics of sniffing bouts may differ based upon what elicited the sniffing. We wanted to know if the sniffing bouts triggered by DA release into the ventral striatum are similar, or distinct from, those which are elicited in response to a stimulus or those bouts spontaneously generated. To address this, we used a machine learning approach (**Fig 5D**). We converted 1s long sniffing traces (the 1s following bout onset) into 1sec long instantaneous sniffing frequency arrays (10ms bin size). Next, we used K-nearest neighbors with Diffusion Time Warping (DTW)^59–61^ to calculate the difference in sniffing frequency dynamics between frequency arrays. With DTW, the smaller the value reported in each time bin corresponds to the dynamics between arrays being more similar at that moment. The model was trained on DTW arrays from odor-evoked, spontaneously-generated, and photostimulation-evoked data (*n*=63-70 arrays/group). Following, we fed the model DTW arrays from each group to test how distinct the sniffing dynamics are between groups. In addition to using the complete ‘real’ data set, we generated synthetic data using SMOTE (synthetic minority oversampling technique^62^) to balance the sample numbers to 70 and separately, tested the model’s classification precision when providing it balanced samples of reduced numbers of 40 or 60 (**Fig 5E**). We found that photostimulation-evoked sniffing was highly distinct from all other types of sniffing. In all runs, precision in classifying opto-evoked sniffing even exceeded the precision of classifying odor-evoked sniffing, which is known to be highly stereotyped (**Figs 5Ei & 5Eii**). As a comparison, spontaneously evoked sniffing was the most poorly classified, likely due to the variability in these self-motivated bouts. While buzz-evoked sniffing was classified better than spontaneous, the model struggled to classify it as well as opto-evoked sniffing. Both SMOTE and the 60 sample runs yielded robust classification of photostimulation-evoked sniffing, with F1 scores (a single metric of model performance) of 0.9 in the 60-sample run, and 1.0 in the SMOTE run. This result implies that DA release into the ventral striatum engages a distinct respiratory pattern from that when an animal engages in self-motivated exploration or upon perception of an environmental stimulus (*i.e.*, an odor).

As an additional experiment, we used an inhibitory optogenetic approach to suppress DAergic terminals in either the NAcSh or the TuS to test whether this achieves reductions in sniffing. We targeted these two ventral striatum subregions based upon their prominent elevations in DA upon sniffing (**Figs 2 & 3**). We injected a Cre-dependent halorhodopsin (AAV5-EF1α-DIO-eNpHR 3.0-EYFP) bilaterally into the VTA of *DAT^IRES^-Cre* or *DAT^IRES^-Cre x Ai9* mice (**Supplementary Fig 7**) and implanted bilateral 400µm diameter optical fibers terminating over the TuS or NAcSh (**Fig 6A & 6B**) to suppress terminal release of DA upon light stimulation. An example of a bilateral cannula implant in the TuS is shown in **Figure 6C**. We sought to understand if suppressing DAergic terminals could reduce sniffing both when mice spontaneously engage in sniffing, and when mice sniff in response to a stimulus. To target spontaneous sniffing bouts, 5s long, 560nm constant light stimulation was delivered through the optical fibers every 20s while mice were in the plethysmograph. Since spontaneous bouts of sniffing can sometimes be short (∼several hundreds of milliseconds), we provided light every 20s without considering if the animal was engaging in a sniff bout or not. We reasoned this would be a more objective stimulation paradigm than if we were to trigger light off of every bout onset (*e.g*., in a closed loop paradigm) which could result in some ‘learning’ of the mouse to change its behavior.

**Figure 6.**
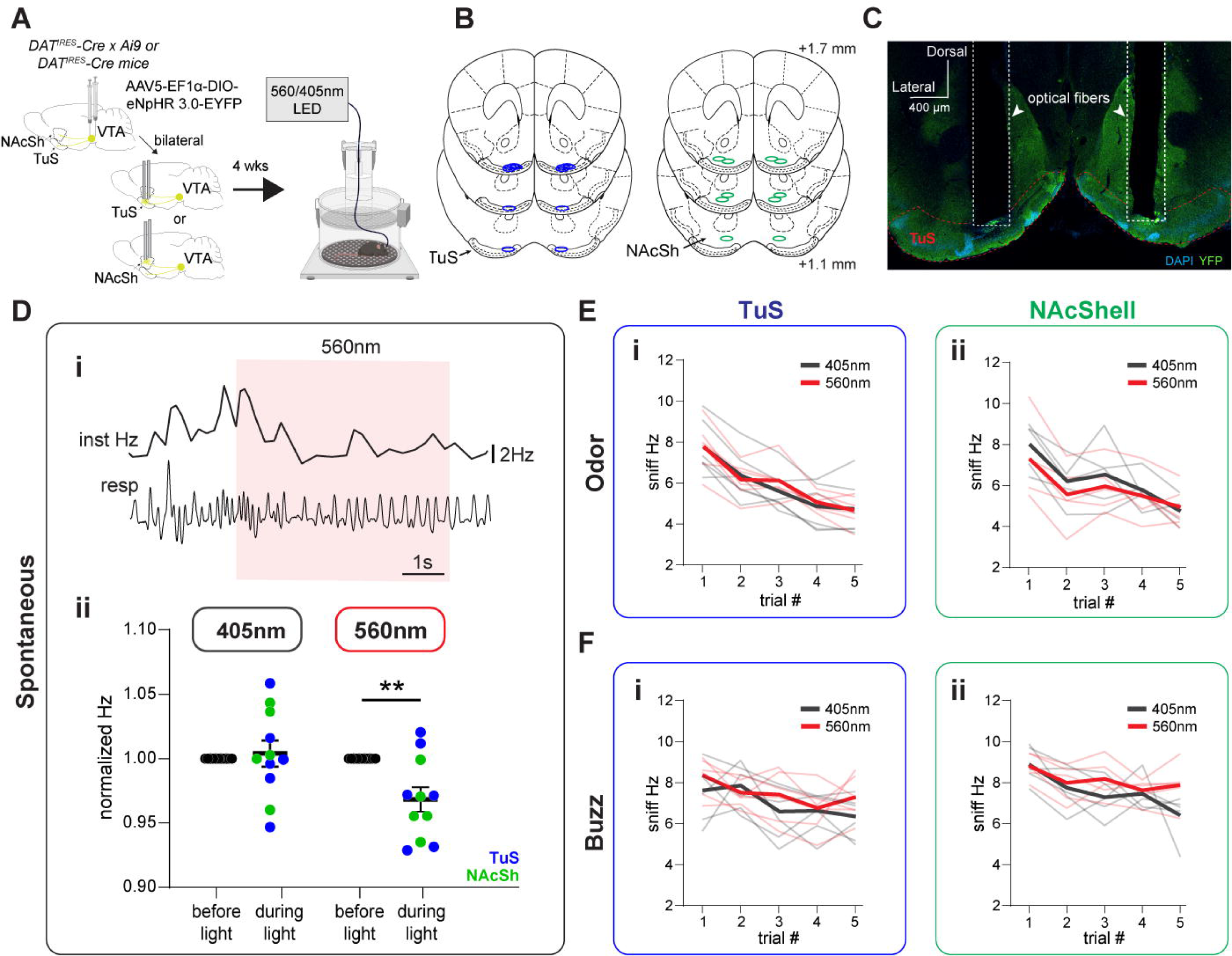
Inhibition of DA release into the ventral striatum and sniffing. A. Schematic of optogenetically-inhibited sniffing paradigm. *DAT^IRES^-Cre* and *DAT^IRES^-Cre x Ai9* mice were injected with AAV-EF1α-DIO-eNpHR 3.0-EYFP bilaterally in the VTA, and bilateral optic fibers terminating the TuS or NAcSh were implanted in the same surgery. At least 4 weeks later, mice were tethered and placed in the plethysmograph to optogenetically inhibit DA terminal release while monitoring sniffing. 560nm light was used to activate eNpHR and 405nm light was used for controls. Image (right) made with BioRender. **B.** Bilateral optic fiber implant locations in the TuS and NAcSh (*n*=5-6/group). **C.** Representative bilateral implant in the TuS. **Di.** Example of respiration during photoinhibition periods. Raw respiratory trace (resp) is shown with interpolated instantaneous frequency above (Inst Hz). 560nm LED duration indicated by red shaded box. **Dii.** Normalized respiratory frequency during the 3s before light stimulation compared to respiratory frequency during 5s light stimulation (405nm stimulation – left, 560nm stimulation – right). Blue dots indicate TuS implanted mice, green dots indicate NAcSh implanted mice. \*\**p*<0.01. **E.** Average instantaneous sniffing frequency of TuS (Ei) and NAcSh (Eii) implanted mice across repeated odor presentations with concurrent delivery of 560nm or 405nm light. Mean during 560nm concurrent light delivery shown in red, with individual mouse data across trials shown in light red. Mean during 405nm concurrent light delivery shown in gray, with individual mouse data across trials shown in light grey. **F.** Average instantaneous sniffing frequency of TuS (Fi) and NacSh (Fii) implanted mice across repeated buzz presentations with concurrent delivery of 560nm (red) or 405nm (gray) light.

As shown in the example in **Figure 6Di**, 560nm light stimulation suppressed sniffing frequency during spontaneous events. Indeed, compared to control mice which received 5s long, 405nm constant light stimulation (light stimulation unable to activate eNpHR^63^), photoinhibition of DAergic terminals in the ventral striatum modestly suppressed respiratory frequency (**Fig 6Dii**; 2-way rmANOVA, effect of light stimulation *F*(1, 10)=11.08, *p*=0.0076; Sidak’s multiple comparisons 560nm before light vs. during light *p*=0.0037). To test effects of DAergic inhibition on stimulus-evoked sniffing, each mouse was presented with pseudorandom trials of odors, including R(+)-limonene, thioglycolic acid, and peanut oil, each five times across a behavior session. 560nm or 405nm light was delivered concurrently with odor and buzz delivery. For this we used a longer light duration (7s, constant) to ensure that light was on upon the possible moment the animal may perceive the stimulus and would typically engage in sniffing. Whereas inhibition of DAergic terminals in the ventral striatum suppressed spontaneous sniffing (**Fig 4D**), there was no effect of photoinhibition on the frequency of odor- or buzz-evoked sniffing in the TuS or NAcSh (**Figs 6E & 6F**, two-way rmANOVA: NAcSh odor no effect of light *F*(1, 4)=1.009, *p*=0.3719; NAcSh buzz no effect of light *F*(1, 4)=1.117, *p*=0.3502; TuS odor no effect of light *F*(1, 5)=0.025, *p*=0.8816; TuS buzz no effect of light *F*(1, 5)=4.696, *p*=0.083). Taken together with the results of the optical stimulation, these results cumulatively support that DA release in the ventral striatum elicits sniffing.

### Ventral striatum neural activity is associated with sniffing

VTA DA is released onto spiny projection neurons in the ventral striatum expressing either the D1 or D2 DA receptor. Is the activity of these neurons associated with sniffing? To test this, we performed two different experiments. First, we took advantage of an existing data set^64^ wherein extracellular neural activity was recorded from the TuS and respiration simultaneously sampled from an intranasal cannula in behaving c57bl/6j mice throughout their engagement in an olfactory discrimination task. In this task, mice were head-fixed which allowed for the snout to remain stable for precision in the odor delivery. A subset of animals contributed respiratory data, and of them, three mice also each contributed multiple single-units (*n*=20 total). As expected from head-fixed animals^65,66^, we observed bouts of sniffing in response to odor delivery and also during the inter-trial interval separating odors (variable 15-17sec window). Further, as expected in head-fixed animals and also seen in mice in the plethysmograph (*e.g.,* **Fig 3**), odor delivery significantly increased the median of the instantaneous respiratory frequency (Wilcoxon rank-sum test, *z*=-19.8, rank sum=3.2×10^7^, *p*<0.0001, **Fig 7C**).

**Fig. 7.**
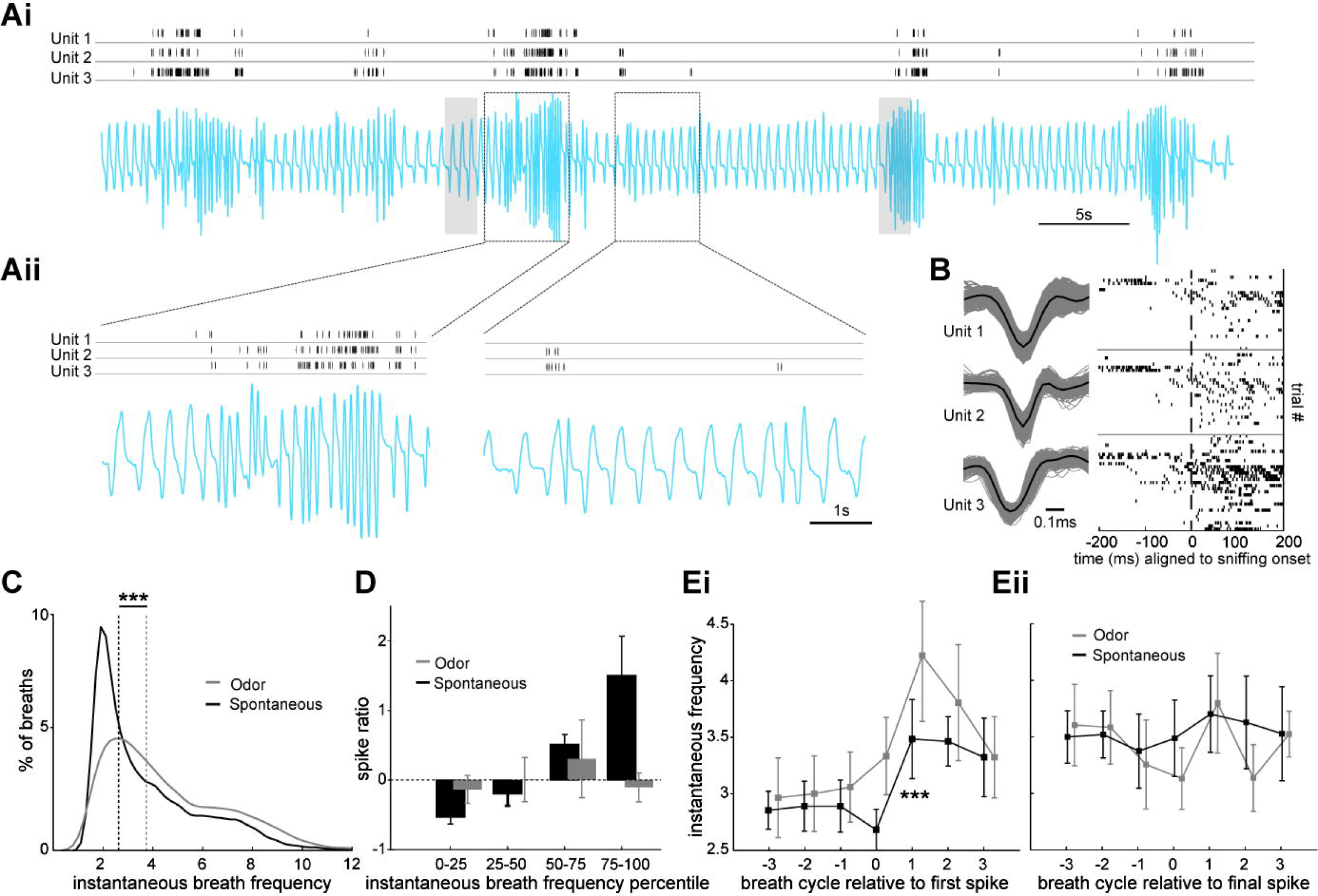
Ventral striatum neuron activity is coupled with sniffing. **Ai.** Continuous recording of intranasal respiration (blue, inhale = upward deflection) along with single-unit activity from three units in the TuS. Grey shaded boxes indicate periods wherein non-novel odors were delivered. **Aii** is a magnified inset to help illustrate the temporal relationship between the three units in **Ai** and sniffing. **B.** Overdrawn waveforms and raster plots of spiking aligned to sniffing onset from the three units in **A** to illustrate that spiking often occurred following sniffing onset. **C.** Distribution of respiratory frequencies from the mice (*n*=3) during both spontaneous and odor-delivery periods (see Methods). **D.** Spike ratio histogram showing that more spiking occurs (*n*=20 units, across 3 mice) during higher frequency respiration than lower frequency respiration, and that this was greater when animals spontaneously transitioned into sniffing during the inter-trial interval than when sniffing during an odor – likely because the different basal respiratory rates in those states (as shown in **C**). **Ei** and **Eii.** Histograms of the firing frequency of the units in D, relative to either the first spike or final spike (Ei or Eii, respectively) indicating that the first spike is predictive that the subsequent breath frequencies will be faster than the breaths prior to the first spike. *** = (ANOVA *F*(6, 112)=4.43, *p*=0.0005; see Results).

As shown in **Figures 7A & B**, we found that the firing of ventral striatum neurons occurs in relation to both bouts of sniffing and even proximal to individual breaths which consisted of a short duration respiratory cycle (*viz*., “sniffs”). Each of the neurons in this example displayed a significant increase in firing in the 200ms following either a sniff or a sniff bout (**Fig 7B**; unit 1 *t*(27)=5.44, *p*<0.0001; unit 2 *t*(27)=2.29, *p*=0.049; unit 3 *t*(27)=-3.46, *p*=0.001). Looking across the entire population of neurons, 19/20 (including the three from **Fig 7B**) were significantly modulated relative to a short respiratory cycle (*p*<0.05, 2-tailed *t*-tests of firing rate 200ms before vs. 200ms during sniffing across sniff occurrences). Among these, they were more likely to fire within a short respiratory cycle during the inter-trial interval when the background respiratory rate is slower (**Fig 7D**, mixed ANOVA, F(3, 76)=31.48, *p*<0.0001).

We then explored the relationship between the probability of a change in firing frequency relative to either the first or last short duration breath in a bout of sniffing. To do this we identified respiration cycles containing an ‘initial spike’ and three respiration cycles before and after that initial spike. We observed that the initial spike occurs before the respiration cycle gets shorter (**Fig 7Ei**). This was especially the case during background breathing in the inter-trial interval when breathing is slower, wherein cycles +1, 2, and 3 (relative to the initial spike) were significantly higher in frequency than cycle 0 (ANOVA *F*(6, 126)=6.37, *p*<0.0001), while cycle 0 did not differ from previous cycles. In the odor presentation period, however, although ANOVA analysis showed there was a significant main effect of respiration cycles (ANOVA *F*(6, 112)=4.43, *p*=0.0005), post hoc comparison showed that cycle 0 was not significantly different from the adjacent cycles. Finally, we performed an additional analysis to determine if the last action potential in a burst was “locked” to the cessation of a sniff bout. **Figure 7Eii** shows that there is no change in respiration cycles aligned to the last spike, whether in the inter-trial interval or odor presentation period. Thus, ventral striatum neural activity appears closely orchestrated with the occurrence of sniffs.

As a second experiment to example the relationship between sniffing and recruitment of post-synaptic ventral striatum neurons, we injected a Cre-dependent AAV expressing GCamP6f (AAV.Syn.Flex.GCaMP6f.WPRE.SV40) into the TuS of D1-Cre and D2-Cre mice^67^ which also were subsequently implanted with a 400µm core optical fiber over the TuS (**Fig 8A & B**). An extra-nasal flow sensor was positioned near the animal’s nose to monitor sniffing^68^. This allowed us to extend the results of the extracellular recordings (though they provided greatest temporal resolution) by analyzing specifically D1 and D2 ventral striatum neuron activity across trials wherein mice spontaneously entered into bouts of sniffing and also during odor-evoked sniffing.

**Figure 8.**
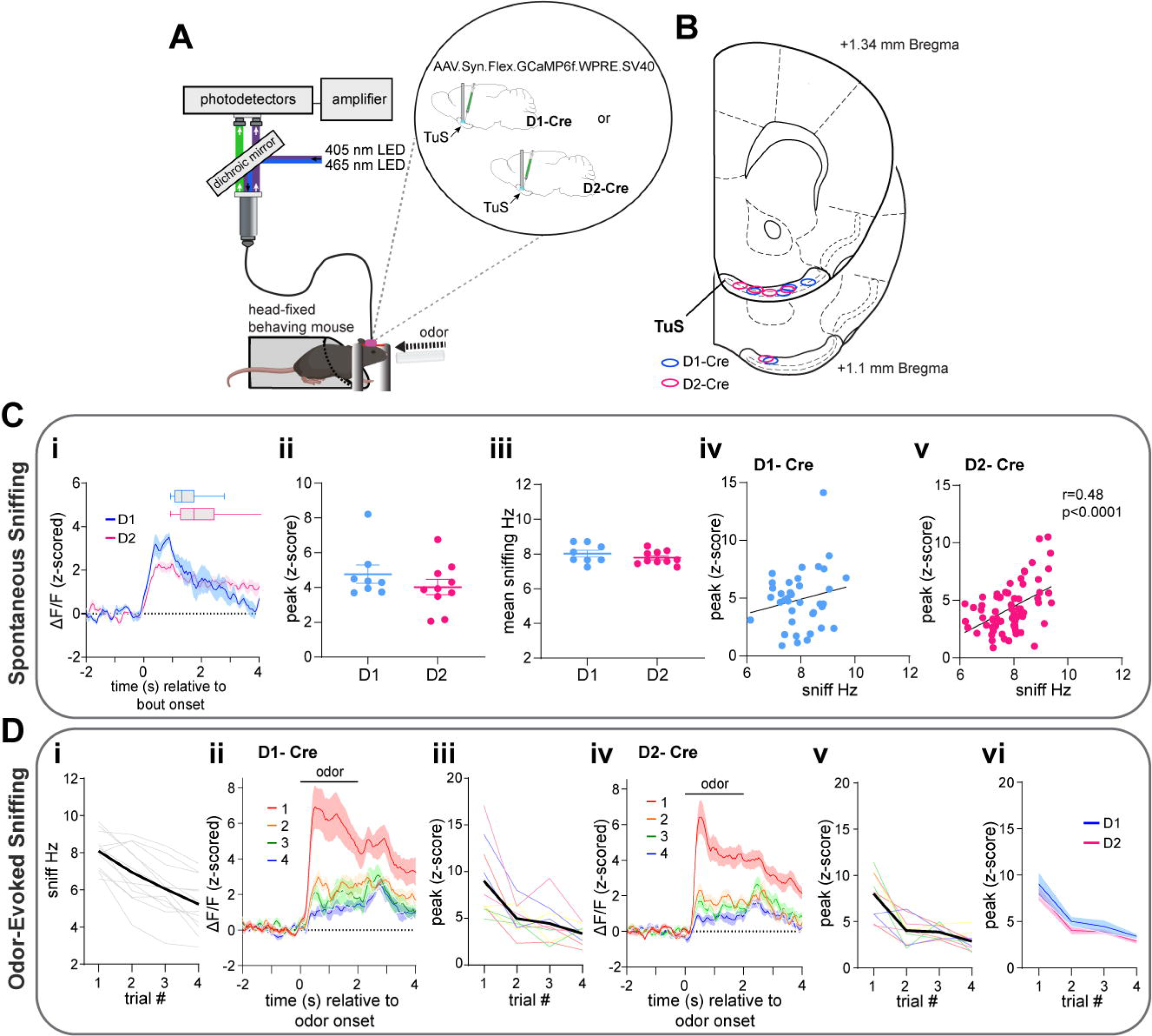
Sniffing is associated with recruitment of D1 and D2 spiny projection neurons. **A.** Schematic of the fiber photometry system for use in head-fixed mice. D1- or D2-cre mice were injected with AAV.hSyn.Flex.GCaMP6f.WPRE.SV40, and implanted with an optical fiber terminating in the TuS. Mice also received a headbar for later fixation. Odors were delivered directly to the animal’s nose, and respiration was recorded with an extranasal flow sensor. Image made with BioRender. **B.** Optic fiber implant locations in the TuS across D1- and D2-cre mice. **C.** GCaMP activity of D1 and D2 neurons in the TuS during spontaneously-evoked sniffing. **Ci.** Averaged z-scored ΔF/F of D1 and D2 neurons aligned to spontaneous sniff bout onset (D1, *n*=4, 2 points/mouse. D2, *n*=5, 2points/mouse. Overlaid box plots represent average bout duration. **Cii.** Average peak of GCaMP evoked z-scored activity during spontaneous sniffing. **Ciii.** Mean sniffing instantaneous frequency during spontaneous bouts. **Civ.** There was no correlation between GCaMP levels in D1 neurons (peak z-score during a bout) during spontaneous sniffing and (average sniff frequency during the bout, *p*>0.05. **Cv.** There was a significant positive correlation between GCaMP levels in D2 neurons during spontaneous bouts and sniffing frequency (*p*<0.0001). **D.** GCaMP activity of D1 and D2 neurons in the TuS during odor-evoked sniffing. **Di.** Average instantaneous sniffing frequency of both D1-cre (*n*=1, 2 sessions/mouse) and D2-cre (*n*=5, 2 sessions/mouse) mice across repeated odor presentations, with individual mouse data across trials shown in light grey. **Dii.** Averaged z-scored ΔF/F across 4 odor presentations in D1-mice. Odor duration depicted by horizontal line. **Diii.** Averaged peak of the z-scored GCaMP response in D1-cre mice across trials. **Div.** Averaged z-scored ΔF/F across 4 odor presentations in D2-mice. Odor duration depicted by horizontal line. **Dv**. Averaged peak of the z-scored GCaMP response in D2-cre mice across trials. **Dvi.** Overlayed averages of peak z-score GCaMP responses from D1- and D2-cre mice across trials. Data in (Ci),(Dii) and (Div) smoothed for visualization purposes only.

Similar to that observed with DA release into the ventral striatum (**Fig 2**), the activity of both D1 and D2 neurons increased upon spontaneous sniffing bouts (**Fig 8C**). D1 and D2 neuron evoked responses were comparable in their temporal dynamics, and both groups displayed similar evoked frequency of sniffing (**Fig 8C**; unpaired *t*(16)=1.084, *p*=0.294). The activity of D2 neurons also positively correlated with the frequency of spontaneous sniffing (**Fig 8Civ & 8Cv**) (D1 Pearson’s *r*(37)=0.19, *p*=0.241; D2 Pearson’s *r*(68)=0.48, *p*<0.0001).

Novel odor delivery (*viz*., the first presentation of a given odor) evoked sniffing in both D1- and D2-Cre mice (paired *t*(11)=11.58, *p*<0.0001), which habituated across subsequent presentations of the same odor (**Fig 8Di**) (rmANOVA *F*(1.73, 19.07)=35.82, *p*<0.0001). During novel odor-evoked sniffing bouts, D1 and D2 neurons increased in their activity (**Fig 8Dii & 8Div)**; D1-Cre: paired *t*(9)=7.03, *p*<0.0001; D2-Cre: paired *t*(9)=9.88, *p*<0.0001). Reminiscent of changes in DA release across repeated trials of odor-evoked sniffing (**Fig 3**), the activity of D1 and D2 neurons similarly reduced across repeated presentations (**Fig 8Diii & 8Dv**; D1-Cre: rmANOVA *F*(1.42, 12.77)=15.69, *p*=0.0008; D2-Cre: rmANOVA *F*(1.80, 16.21)=25.01 *p*<0.0001). The evoked neural activity and its subsequent adaptation across trials was similar in D1- and D2 mice (**Fig 8vi**; two-way ANOVA, no effect of genotype *F*(1, 18)=1.387, *p*=0.254). These results indicate that the activity of DA receptor-expressing ventral striatum neurons post-synaptic from midbrain DAergic terminals is associated with sniffing.

### DA receptor antagonism in the ventral striatum reduces both the occurrence and frequency of sniffing

Ventral striatum neural activity can be influenced by a wealth of neurotransmitters/neuromodulators arising from numerous bottom-up and top-down brain regions. For instance, NAc activity is influenced not only by DA, but also by glutamatergic inputs from the frontal cortex^69^. While the above results suggest that sniff-evoked changes in mesolimbic DA influx to the ventral striatum engages postsynaptic neurons, those results still beg the question: is DA binding to DA receptors on ventral striatum neurons needed for an animal to initiate sniffing?

We implanted c57bl/6j mice of both sexes with bilateral indwelling infusion cannulae into either the NAcSh or TuS (**Fig 9A**). We focused on antagonism of DA receptors in the TuS and NAcSh since it was within these regions wherein we found the strongest correlation between DA levels and sniffing frequency (**Figs 3 & 4**). Since TuS and NAcSh neurons may express D1, D2, or D3 receptors^70–74^, across days in a counter-balanced within subject’s design, each mouse was intracranially infused with SCH23390, raclopride, or PG01037 to selectively inhibit D1, D2, or D3 receptors, respectively. The infusion occurred 10min prior to placing the mouse into the plethysmograph for respiratory monitoring, wherein we measured both spontaneous and sensory-evoked sniffing bouts (**Fig 9B)**. We analyzed for effects of D1, D2, and D3 antagonism on both the number of sniff bouts and their frequency, displayed spontaneously and stimulus-evoked. DA receptor antagonism has some known sedative effects^75^, and notably, DA receptor antagonism had some effect on baseline breathing, with specifically D2 receptor antagonism in the NAcSh influencing breath frequency (Wilcoxon signed-rank test *W*=-279, *p* =0.012; **Supplementary Fig 3**).

**Figure 9.**
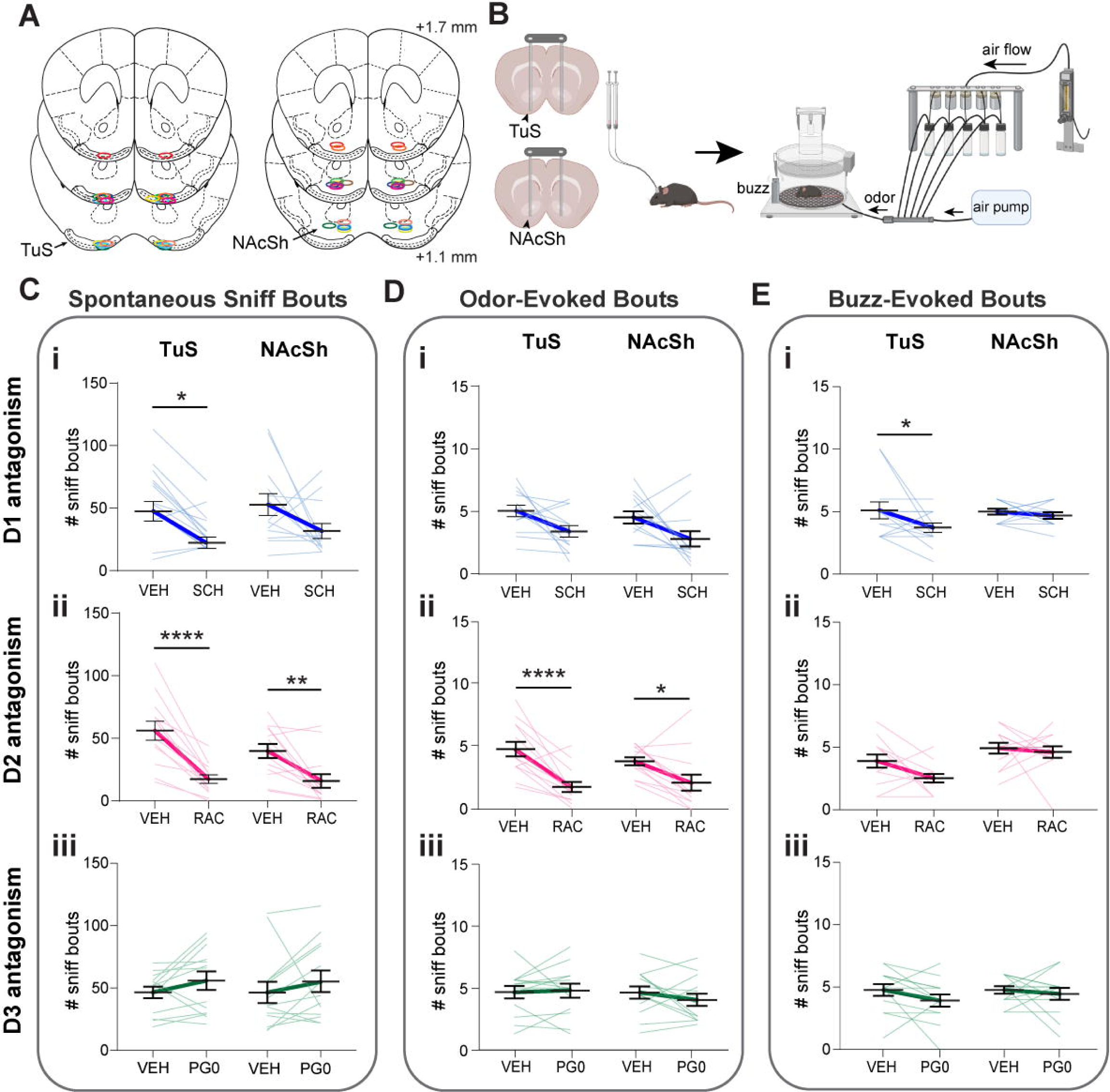
Dopamine binding to D1 and D2 ventral striatum receptors contributes to sniffing. A. Bilateral cannulae implant locations in the TuS and NAcSh. Colored for visualization purposes only. Position relative to bregma. B. Diagram of pharmacological manipulation and behavioral paradigm. Animals were implanted with stainless steel, bilateral cannulae with tips terminating in the TuS (*n*=13) or NAcSh (*n*=13). After surgical recovery, animals underwent behavioral testing. In a counterbalanced, within-subjects design, animals were administered an intracranial infusion of a selective D1, D2, or D3 antagonist or corresponding vehicle control 10min prior to being placed in the plethysmograph. Vehicle = sterile saline + 0-12% DMSO. During a behavioral session, animals were presented with both olfactory and non-olfactory (buzz) stimuli. Image made with BioRender. **C.** Impact of selective antagonism of D1-(Ci), D2-(Cii), and D3-receptors (Ciii) on spontaneous sniffing. D3 antagonism had no effect on spontaneous sniffing in the TuS or NAcSh (Ciii, *p*>0.05). **D.** Impact of selective antagonism of D1-(Di), D2-(Dii), and D3-receptors (Diii) on odor-evoked sniffing. D1 and D3 antagonism had no effect on odor-evoked sniffing in the TuS or NAc (Di & Diii, *p*>0.05). **E.** Impact of selective antagonism of D1-(Ei), D2-(Eii), and D3-receptors (Eiii) on buzz-evoked sniffing. D2 and D3 antagonism had no effect on buzz-evoked sniffing in the TuS or NAc (Eii & Eiii, *p*>0.05). Data are mean (dark bolded line) ± SEM, lighter lines = individual mice. Abbreviations: SCH (D1 antagonist SCH23390), RAC (D2 antagonist raclopride), PG0 (D3 antagonist PG010370), VEH (vehicle). \*\*\*\**p*<0.0001, \*\**p*<0.01, \**p*<0.05.

We found that D1 receptor binding contributed to spontaneous sniffing in that D1 receptor antagonism in the TuS reduced the number of spontaneous sniffing bouts (**Fig 9Ci**; two-way ANOVA, main effect of treatment *F*(1, 24)=12.62, *p*=0.002; Sidak’s multiple comparisons TuS VEH vs. SCH *p*=0.023). D1 receptor binding also contributed to sensory-evoked sniffing in that D1 receptor antagonism in the TuS reduced the number of buzz-evoked bouts (**Fig 9Ei**; two-way ANOVA, main effect of treatment *F*(1, 24)=4.30, *p*=0.049; Sidak’s multiple comparisons TuS VEH vs. SCH *p*=0.048).

Likewise, we found that D2 receptor binding contributed to spontaneous sniffing in that D2 receptor antagonism in both the TuS and NAc reduced the number and frequency of spontaneous sniffing bouts (**Fig 9Cii**; two-way rmANOVA, main effect of treatment *F*(1, 24)=48.64, *p*<0.0001; Sidak’s multiple comparisons TuS VEH vs. RAC *p*<0.0001, NAc VEH vs. RAC *p*=0.002). D2 receptor binding in the TuS and NAc also contributed to odor-evoked sniffing in that D2 receptor antagonism reduced the number of sniffing bouts (**Fig 9Dii**; two-way rmANOVA, main effect of treatment *F*(1, 24)=31.26, *p*<0.0001; Sidak’s multiple comparisons TuS VEH vs. RAC *p*<0.0001, NAc VEH vs. RAC *p*=0.017). In contrast, we saw no effect of D3 receptor inhibition on spontaneous or odor-evoked sniffing in either the TuS or NAcSh (**Fig 9Ciii, 9Diii, 9Eiii**).

Interestingly, D2 antagonism in both the TuS and NAc not only reduced the number of odor-evoked bouts displayed by mice, but in both regions, the odor-evoked bouts which were displayed were reduced in frequency (**Supplementary Fig 8Bi & 8Bii**; D1 two-way rmANOVA main effect of treatment *F*(1, 24)=7.75, *p*=0.010; Sidak’s multiple comparisons TuS VEH vs. SCH *p*=0.228, NAcSh VEH vs. SCH *p*=0.056; D2 two-way rmANOVA, main effect of treatment *F*(1, 24)=96.64, *p*<0.0001; Sidak’s multiple comparisons TuS VEH vs. RAC *p*<0.0001, NAcSh VEH vs. RAC *p*<0.0001). This was not observed during spontaneous sniffing bouts or buzz-evoked sniffing bouts pointing to a unique system which supports the frequency of odor-evoked sniffing compared to spontaneous sniffing which was influenced solely in terms of the total number of bouts. Altogether, these results confirm that mesolimbic DA initiates sniffing by its actions upon D1 and D2 receptors in the TuS and NAcSh.

## Discussion

DA appears to partake in numerous functions^33,34,76,77^. For instance, DA can influence the perceived salience of stimuli wherein DA might make certain smells stand out more^39^, making them more noticeable during sniffing. DA might enhance our sensitivity to sensory stimuli, including odors, allowing them to be better detected at low levels^78^. DA is also involved in arousal and attention mechanisms wherein it could heighten our alertness and focus on sensory inputs, including odors and other cues^79,80^. Indeed, DA supports cognitive functions in humans during non-olfactory functions such as during gazing^81^. Further, DA is involved in synaptic plasticity and learning from experiences and adapting to new information. When sniffing odors, DA release, including upon ventral striatum neurons, could contribute to the brain’s ability to form associations and learn information^30^. Likewise, when we encounter pleasant or novel odors, DA release might contribute to feelings of pleasure and satisfaction^53^. Indeed, DA is a key transmitter in the brain’s reward system and is connected to motivation and exploration^82^, wherein it may influence our motivation to sniff, encouraging us to explore and investigate different odor landscapes. Furthermore, several lines of evidence have documented the orchestration of sniffing during motivated behaviors and even upon anticipation of a reinforcer. These functions of DA stated, no prior work has mechanistically linked DA with the highly conserved and ethologically-relevant behavior of sniffing. Our results support a model wherein DA acts upon the ventral striatum, specifically upon neurons with D1 and D2 receptors in the TuS and NAcSh, to initiate sniffing and even increase the frequency of sniffing. This finding perhaps is analogous to the known role of DA in guiding attention in other sensory modalities, including in humans^83,84^ We predict the promotion of sniffing by DA is an essential component linking the historic cooccurrence of sniffing with motivated states.

Stimulus active sampling behaviors have a tight linkage to both the acquisition and processing of environmental information, and these are modulated by ascending monoaminergic neurotransmission. For instance, stimulation of the locus coeruleus in rats increases the responsiveness of barrel cortex neurons to whisker deflection^85^. In zebra finches, auditory coding in the finch auditory cortex homologue is modulated by norepinephrine^86^. Likewise, visual processing in the lateral geniculate nucleus of cats is modulated by dopamine^87^. In terms of active sampling behaviors and their regulation by monoaminergic transmission, rhythmic licking in mice can be suppressed by DA D1 receptor antagonism in the motor cortex^88^. The rhythmic movement of the vibrissae (“whisking”) is perhaps the most well-studied in terms of its modulation by monoaminergic inputs. Serotonergic input from the dorsal raphe nucleus, whose neurons fire rhythmically within the whisking frequency bands, generates whisking in rats and whisking can be diminished by serotonin receptor antagonism in the dorsal raphe^89^. Also, recent evidence indicates that mesolimbic DA input to the NAc in mice promotes orofacial behaviors including whisking^41^. Our results add to these prior studies by establishing a necessary and sufficient role for mesolimbic DA in the initiation of sniffing behavior in that not only does photostimulation of DAergic terminals initiate sniffing bouts, but also, blockade of DAergic receptors and optical inhibition of DAergic terminals suppresses the normal occurrence of sniffing and its frequency. In line with these results, VTA→ventral striatum DA may serve as a pathway to facilitate adaptive sniffing to promote olfactory search behavior, and moreover may shape odor processing given the regulation of odor coding and perception by the respiratory rhythm^90–92^. Given the extensive innervation of the olfactory bulb by neuromodulatory systems, including norepinephrine, and the presence of DAergic interneurons in the olfactory bulb, one might speculate that odor processing and perception are resultant from a delicate balance of neuromodulators^93^, which not only recognize a novel odor, but also consequentially instruct novel motor patterns. The time-course of stimulation-evoked sniffing we report herein supports that indeed DAergic release into the ventral striatum may not be the initial driver of novel-odor evoked sniffing, which can be as fast as ∼100ms^94^.

We found heterogeneity of DA release in ventral striatum subregions. Previous work from Menegas and colleagues reported that DA is only released in the tail of the striatum upon novel odor presentation (which likely would entail sniffing), with negligible changes in DA levels seen in the ventral striatum^32^. It is interesting to note that in their study, ventral striatum recordings were largely acquired from the NAcC, wherein we similarly did not see increases in DA levels. Furthermore, the results of Menegas and colleagues even suggest subtle reductions in DA levels, which agrees with our current findings from the NAcC. We expand upon this work by investigating DA dynamics across all three ventral striatal regions which include the NAcC, NAcSh, and TuS while animals sniff. Our findings are reminiscent of those monitoring DA release during other behaviors which has uncovered heterogeneous levels of DA in the NAcC versus NacSh in manners likely supported by differing VTA neural pathways innervating these subregions^43,44,95^. Here we found that in comparison to the increases in DA upon sniffing in both the TuS and NAcSh, that the NAcC experienced reductions in DA release (**Figs 2 & 3**). This implies that sniffing increases tonic firing of VTA DAergic neurons, or suppresses firing of DAergic neurons, in the NAcSh and TuS and NAcC, respectively. That optical excitation of the terminals in any of the regions equally initiates sniffing suggests that this is overriding changes in VTA neuron firing which give rise to DA dynamics in the ventral striatum and also hints that heterogeneity in subregion DA dynamics are not a byproduct of interregional differences in DA clearance (*e.g.,* by different kinetics or function of the dopamine transporter). It is tempting to speculate that different DA dynamics in these subregions might stem from unique inputs from areas that may signal salience or arousal which would synapse on VTA GABAergic neurons, including projections from the TuS and NAc themselves^95^. Furthermore, different DA neuron populations in the VTA display heterogenous responses to novelty vs. reward^32,80,96,97^, which might stem from functional subtypes of DAergic neurons. It will be important to investigate the differing VTA neuron populations which may underly these subregion-specific differences in DA dynamics in the ventral striatum during sniffing.

What ‘type’ of sniffing does mesolimbic DA initiate? The machine learning model trained on different experimenter classified sniffing patterns (**Fig 5D**) points to mesolimbic DA-evoked sniffing as its own type of sniffing, unique from that a mouse would self-motivate upon the desire to explore (spontaneous) and that evoked by a sensory stimulus (like an odor or buzz). So, what type is it? Sniffing is a dynamic behavior which can be displayed in unique manners based upon context. For instance, dogs sniff at lower frequencies when searching the air for odors than when searching on the ground wherein they elevate their sniffing rate^98^. Rats also display sniffing in a distinct frequency band when transitioning from sampling odors to anticipating a reinforcer in an instrumental task in manners which suggests sniffing reflects strategic ‘modes’ or states^5^. Our results are in-line with these prior works and suggest that mesolimbic DA-evoked sniffing reflects a unique sniffing mode. Several reports have established links between specific brain regions, and neurons within those brain regions, in regulating adaptive respiration, although the role of those regions does not appear to initiate sniffing as much as shape more specific respiratory parameters (*e.g.* ^99^). Our results wherein we varied photostimulation of DAergic terminals uncovered that the duration of evoked-sniffing bouts did not increase with more prolonged optical stimulation (**Fig 4H**). Considering that VTA DAergic neurons fire in characteristic phasic manners^100,101^, this suggests that DA release into the ventral striatum is more so to initiate and possibly elevate sniffing frequency than to maintain its tonic display.

There are a few caveats to this study we believe important to note. First, optogenetic stimulation of DAergic terminals, while herein an effective means at robustly driving sniffing, is likely driving DA release onto postsynaptic neurons in supraphysiological manners. While we selected a stimulation paradigm established by others to be effective on TuS projecting VTA terminals, it is nevertheless likely DA is released at higher than usual levels. Nevertheless, both our pharmacological data (**Fig 9**) and our work wherein we removed *VGluT* from tyrosine hydroxylase expressing neurons (**Supplementary Fig 6**) all agree that DA is contributing to the initiation of sniffing. It is also possible that some behavioral effects from the photostimulation are due to backpropagation of activity into the VTA itself. Again though, the pharmacological results indicate DA input to the ventral striatum is important for sniffing. Second, while DA release into the ventral striatum initiates sniffing, and while in several of our analyses, DA levels were correlated with sniffing frequency, this was not always the case. For instance, for the buzz stimulus experiments, large DA levels were found despite the mice only displaying modest frequencies of sniffing. This corresponds with other studies which have reported DA release in response to a variety of salient stimuli. Finally, while D2 receptors are enriched on spiny projection neurons in the TuS and NAc, they also serve autoreceptor functions on VTA DAergic terminals, including those innervating these regions. So, while we know local D1 antagonism suppressed sniffing, it is possible the effects of D2 antagonism are due to contributions of antagonizing D2 receptors on both the VTA terminals (which would enhance DA release) and on spiny projection neurons.

Antagonizing DAergic receptors versus inhibiting DAergic terminals resulted in different effects upon respiration. Whereas pharmacological inhibition of TuS D1 receptors and NAc D1 and D2 receptors influenced specific occurrences of sniffing, including both spontaneous and stimulus-evoked sniffing (**Fig 9**), optical inhibition of DAergic terminals only impacted spontaneous events (**Fig 6**). We reason this may be due to a few possibilities. In our photoinhibition experiments, we targeted the TuS and NAcSh separately. Given the transient and specific nature of the photoinhibition paradigm, it is likely that the inhibition of DA release in the TuS could not override the resultant effects of DA release in the NAcSh on sniffing, or *vice versa*, therefore resulting in modest changes in respiration. Why then did we see robust effects when we pharmacologically inhibited DA receptors in a single region (**Fig 9 & Supplementary Fig 8)**? We reason this may be because a direct antagonist infusion would inhibit a great number of post-synaptic receptors, and this coupled with potential influences of the antagonists on autoreceptor function and the massive collateralization of spiny projection neurons upon each other would yield more dramatic changes in the ability of the cells to act upon presynaptic DA.

A major question is how the receipt of DA upon spiny projection neurons might ultimately drive changes in the respiratory rhythm. Both the TuS and NAc have outputs into downstream structures important for homeostatic regulation/control and limbic function. Ventral striatum spiny projection neurons also feedback upon the VTA in manners regulating DA release and this too could support, even if just bisynaptically, modulation of respiratory central pattern generators which are sensitive to dopamine^102,103^. Perhaps one clue may reside from our results wherein we attempted to stimulate sniffing under anesthesia and found that DA release into the TuS or NAc is incapable of initiating sniffing (**Fig 5A & B**). This suggests that the functional ‘sniffing’ output from the ventral striatum is within a structure which can drive changes in breathing but not under anesthesia. It is not uncommon for stimulation of some respiratory control centers to be incapable of altering respiration under anesthesia^21,103^. For instance, the ventral striatum projects to hypothalamic nuclei that are known to innervate respiratory control nuclei in the respiratory pons and could support entrainment of breathing at sniffing frequencies. Interestingly, these hypothalamic targets are also state dependent in that changes in breathing frequency do not occur upon stimulation under anesthesia ^21^. Thus, there seems to be somewhat of a paradox in that in some respiratory structures, including the PreBӧtzinger complex, high frequency stimulation is incapable of driving respiration at frequencies greater than 6Hz^104,105^. Identifying what downstream circuits link the TuS and NAc to respiratory brain stem circuits to orchestrate sniffing remains an important goal. Our results point to D1 and D2 ventral striatum neurons being part of a feedforward/feedback loop between the basal ganglia and respiratory control regions with DAergic VTA neurons being an intermediary circuit node. While the bio-sensor based fiber photometry recordings we used to capture changes in DA and ventral striatum D1 and D2 neurons suggest that these signals may at times lag the onset of a sniff bout, we did not correct for the lag inherit in photometry recordings. Despite that, in several of the examples provided, DA levels begin to rise immediately upon sniff bout onset (*e.g.*, **Fig 2Eii**). Further, the TuS single unit recordings offered temporal dynamics which point to firing of at least some TuS neurons immediately prior to the detectable increase in sniffing. This ultimately leads us to speculate that a signal arrives at the ventral striatum (regarding a novel odor, or a change in environment or internal state) which is ultimately routed through basal ganglia to initiate sniffing by means of actions in the respiratory brainstem. At that point, DA is released by activation of VTA DAergic neurons (perhaps as even modulated by ventral striatum input itself, e.g.,^106,107^), which then, based on our results, would further drive (“invigorate”) sniffing by its actions on those same ventral striatum neurons. If future studies support this reciprocal DAergic loop model for sniffing, naturally they would need to incorporate inputs/outputs to frontal cortices, hypothalamic structures, motor cortex, and other regions integral for the plethora of behaviors and states which might coincide with sniffing. While in many of our measures the frequency of sniffing positively correlated with the peak magnitude of DA levels (*e.g.,* **Figs 2 & 3**), some instances of sniffing did not co-occur with major rises in DA (*e.g.,* **Supplementary Fig 2 and also as shown in Figs 2Fii & 2Gii**) and so it is likely multiple circuits are at play which link DA with sniffing.

In conclusion, we report evidence that mesolimbic DA input into the ventral striatum contributes to the display of the widely conserved behavior of sniffing. The nature of sniffing being integral to both olfaction and motivated behaviors implicates this circuit in a wide array of functions. Especially in macrosmatic animals, like the mice studied herein, olfaction is crucial for appropriate social behaviors, avoiding predation, and foraging. Based upon our evidence, we propose it is likely this circuit provides a neuromodulatory means whereby motivated states signaled by DA release tune the occurrence and frequency of sniffing in order for animals to thrive in the above ethological scenarios.

## Methods

### Animals

Male and female mice were housed on a 12:12hr light-dark cycle with *ad libitum* access to food and water. All behavioral testing took place during the light cycle. Mice with viral injections alone were group housed (≤5 mice/cage) and any mouse with a chronic implant was single housed following its surgical procedure. Food (Envigo Teklad Global, 18% rodent diet irradiated pellet, Cat # 2018; Indianapolis, IN) and reverse osmosis water were available *ad libitum* except during behavioral monitoring/recordings. Mice were housed in standard shoebox sized plastic cages (Allentown Jag 75 micro-vent system; Allentown, PA; L: 29.2cm W: 18.5cm H:12.7cm) on a rack made for individually vented cages. Corncob bedding and a Nestlet^TM^ (Ancare; Bellmore, NY) were in each cage along with manzanita sticks (Bio-Serve). Mice were housed on a 600-1800hr light cycle with lights on during the daytime. Temperature in the room averaged 70 ± 2°F, with 30-70% humidity and 10-15 room air changes per hour. The housing room was specific pathogen free. Observation occurred from 1000-1700hrs. All animal care was conducted with in the AAALAC animal research program of the University of Florida, in accordance with the guidelines from the Guide for the Care and Use of Laboratory Animals and approved by the University of Florida Institutional Animal Care and Use Committee.

Mouse lines included the following transgenic lines which all were maintained on a c57bl/6j background (Jackson labs, strain #000664; RRID:IMSR_JAX:000664) and were bred in house within a University of Florida vivarium. *drd1-Cre* and *drd-2-Cre* lines^67^ were obtained from the UC Davis Mutant Mouse Regional Resource Center (D1: EY262Gsat/Mmucd; D2: ER44Gsat). *DAT^IRES^-Cre* mice^108^ (strain #006660; RRID: IMSR_JAX:006660), *TH-Cre* mice^109^; strain #008601; RRID: IMSR_JAX:008601), *VGluT2 fl/fl mice*^110^; strain #012898; RRID: IMSR_JAX:012898), and *Ai9* TdTomato Cre reporter mice^111^ (strain # 007909, IMSR_JAX:007909) were obtained from Jackson labs. A total of 178 mice (mixed sexes) were used in this study for experimentation. Out of these, a total number of 51 were excluded from data analyses due to experimental issues (*e.g.,* surgical complications, off-target brain implants, viral injections which missed the target).

For all surgical procedures, under aseptic conditions, mice were anesthetized with Isoflurane (2-4% in oxygen, IsoFlo®, Patterson Veterinary) and mounted in a stereotaxic frame where they were maintained under Isoflurane. Body temperatures were maintained with a 38°C heating pad underneath. A local anesthetic, bupivacaine hydrochloride (Marcaine™, 5mg/kg subcutaneously, s.c., Patterson Veterinary), and the analgesic meloxicam (5mg/kg, s.c., Patterson Veterinary) were administered before incision and exposure of the skull. Mice receiving chronic implants (*e.g.,* cannulae, optic fibers) were given meloxicam (0.5mg/kg, s.c.) for 3 days following surgery.

For viral injections, craniotomies were made over the ventral tegmental area (VTA) (−3.6AP/0.35ML/-4.75DV), tubular striatum (TuS) (1.4AP/1ML/-4.90DV), nucleus accumbens shell (NAcSh) (1.4AP/1ML/-4.40DV), and/or nucleus accumbens core (NAcC),(1.4AP/1ML/-3.90DV) and a pulled glass micropipette containing the AAV was lowered into the region of interest. Viruses were all delivered via the pulled pipette attached to a Nanoject III (Drummond Scientific) at a rate of 2nl/s unless otherwise noted. After waiting 10min, the pipette was slowly withdrawn from the brain. Unless followed by an implant in the same surgery, craniotomies were sealed with dental wax, and the incision closed with wound clips.

For intranasal ‘sniff’ cannula implantation, a ∼1mm diameter hole was drilled through the right nasal bone, 1.5mm anterior the frontal/nasal fissure and 0.5mm lateral following prior methods ^6^. A stainless-steel guide cannula (22GA, #C313G, P1 Technologies, Roanoke, VA) cut to extend 0.5mm below the pedestal was lowered through the craniotomy and secured with dental cement. A dummy cannula (#C313DC, P1 Technologies, Roanoke, VA) cut to extend 0.2mm past the guide cannula was also inserted when not recording from the mouse.

To record respiration, the dummy cannula was removed and a pressure transducer was connected (CPXL04GF, Honeywell) to the cannula via a flexible piece of polyethylene tubing. The mouse was then placed in the plethysmograph and recorded for 15min. Plethysmograph-recorded transients were detected using a flow transducer (Data Sciences International) and digitized at 300Hz (0.1-12Hz band pass; Tucker Davis Technologies, Alachua, FL), following a 500x gain amplification (Cyngus Technology Inc, Southport, NC). Intranasal pressure transients were amplified and digitized using the same settings.

### Anatomical Tracing

For synaptophysin injections, a craniotomy was made over the VTA, and a micropipette containing 100nL of pAAV-hSyn-FLEx-mGFP-2A-Synaptophysin-mRuby (Addgene 71760-AAV1; titer: 9.8×10^12^vg/mL) was injected unilaterally into the VTA of *DAT^IRES^-Cre* mice.

3 weeks following injections, mice were overdosed with sodium pentobarbital (Fatal-Plus, Patterson Veterinary) and transcardially perfused with 0.9% saline followed by 10% phosphate-buffered formalin. Brains were removed and stored in a 10% formalin in 30% sucrose solution at 4°C prior to sectioning. Brains were coronally sectioned at 40µm and sections were stored in Tris-buffered saline with 0.03% sodium azide.

#### Image acquisition and quantification

Brain regions of interest were identified using the mouse brain atlas^112^. For the TuS, 12 images were acquired spanning 1.94mm - 0.14mm anterior to Bregma. For the nucleus accumbens, a total of 8 images were acquired spanning 1.94mm – 0.74mm anterior Bregma.

Images were acquired with a Nikon Eclipse Ti2e fluorescent microscope. For brains with synaptophysin injections (*n*=6 mice), images were acquired at 20x magnification across the hemisphere ipsilateral to the injection site, and Z-stacked at every 4μm. For quantification, regions of interest (ROIs) were drawn around areas of interest, namely the NAcC and NAcSh along with the medial and lateral TuS. To determine the medial and lateral portions, the TuS was divided into thirds and the most medial or lateral third was used to ensure distinctness between the two regions^113^. Images were preprocessed to decrease average background fluorescence and enhance contrast. A semiautomated counting algorithm created within NIS elements software (Nikon) was used to detect and count fluorescent puncta, allowing for an unbiased estimation of puncta numbers. Puncta were identified and quantified based on their size and fluorescence intensity via bright-spot detection.

To reconstruct synaptophysin injection sites **(Supplementary Fig 1)**, the VTA was imaged from −2.92 to −3.80mm posterior bregma, as identified with use of a brain atlas^112^. Next, an experimenter (A. C.-L.) identified and marked GFP+ cell bodies in VTA schematics within Adobe Illustrator.

### Fiber photometric recording in the plethysmograph

#### Virus injection and optical fiber implant

C57bl/6j mice were injected unilaterally in the NAcSh, NAcC, or TuS (*n*=7/group) with 100nL pAAV.hSyn-GRAB_DA1h (Addgene 113050, titer: 1×10^13^vg/mL)^50^. For controls, c57bl/6j mice were injected unilaterally in NAcSh, NAcC, or TuS (*n*=2-4/group) with 100nL pAAV.hSyn-GRAB_DA-mut (Addgene 140555, titer: 1×10^13^ vg/mL). Viral injections were performed as previously described (see ‘Animals’). In the same surgery, a 400µm core, 0.48NA optical fiber secured in a 2.5mm outer diameter metal ferrule was chronically implanted to terminate in the TuS, NAcSh, or NAcC and affixed to the skull with dental cement. Animals were recorded from 3-5 weeks following surgery.

#### Photometry

465nm (GRAB_DA_/GRAB_DA_-mut excitation wavelength, driven at 210Hz) and 405nm (UV excitation wavelength, driven at 330Hz; control) light emitting diodes (TDT, RZ10x) were coupled to a fluorescent minicube (Doric lenses, FMC6) via 200µm, 0.22NA FCM optic fiber patch cords. Both excitation and emission light were directed through a single 400µm, 0.57NA patch cord connected to the animal by the implanted 2.5mm metal ferrule. Emission light was directed through the mini cube and coupled to femtowatt photoreceivers to monitor GFP and UV signals (TDT, RZ10x). The emission signals were low-pass filtered (6^th^ order), pulse demodulated, and digitized (1kHz) using a Tucker Davis Technologies processor (TDT). All parameters for imaging and acquisition of photometry data were consistent across all groups.

#### Respiratory data acquisition

To simultaneously monitor photometry signals and respiration, mice were gently tethered to the patch cord and placed in a whole-body plethysmograph (Data Sciences International, St. Paul, MN). Respiratory transients were detected using a flow transducer (Data Sciences International) and digitized at 300Hz (0.1-12Hz band pass; Tucker Davis Technologies, Alachua, FL), following a 500x gain amplification (Cyngus Technology Inc, Southport, NC). Within all plethysmography recordings, we analyzed the instantaneous frequencies of the peak-detected respiratory signal to identify periods indicative of sniffing^6^. Since mice rapidly transition in and out of sniffing bouts ^6^, we restricted our analyses to more stable/pronounced bouts versus short bouts by only defining a sniff bout as the occurrence of sniffing ≥6Hz for ≥1s in duration. Further, in some instances, mice continuously sniff for numerous seconds and then reengage in another bout shortly thereafter. Since these may reflect periods of more generalized arousal versus spontaneous exploratory behavior, we excluded sniff bouts occurring in proximity to these long (>10s) sniffing bouts. In a subset of mice, an observer *post-hoc* monitored video (10Hz, 640×480 resolution) which was recorded simultaneously with respiratory signal from the plethysmograph to validate that focusing on >6Hz changes in respiration was indicative of ‘active sampling’ behaviors from the mouse (including snout directed investigation and head-turning) and found 1:1 correspondence supporting use of respiratory signals from the plethysmograph as suitable for detecting active investigation (of course in this set-up we could not see minor facial, motor or vibrissal movements).

#### Behavioral monitoring

Behavioral sessions lasted approximately 30min during which mice were presented with both olfactory and non-olfactory stimuli. To allow for examination of spontaneous sniffing, we included 5min periods where no stimuli were presented at the beginning (0-5min), middle (12-17min), and end (28-33min) of a session. Spontaneous bouts for analysis were identified between 2-5min, 14-17min, and 30-33min to ensure that they were far removed from any stimuli presentations. Between 5-12min, 2 trials of 4 different stimuli were pseudorandomly presented. The final 3 trials of pseudorandom stimuli presentations occurred between 17-28min. Each stimulus was presented for 2s with a 60s inter-trial interval.

#### Stimulus delivery

Odors R(+)-limonene and thioglycolic acid (Sigma-Aldrich, >97% purity), and that of peanut oil (La Tourangelle®, Woodland, CA), were presented to mice via a custom air-dilution odor delivery device connected to the plethysmograph (see Results for rationale in selecting these specific odors). During experimentation, 2mL of all odors were contained in 40mL glass headspace vials at room temperature. Computer-controlled and user-initiated openings of valves allowed for odors to pass through headspace vials at a flow rate of 50mL/min, following which they were mixed with a carrier stream of clean filtered room air delivered by a pump (1L/min; Tetra Whisper, Melle, Germany) before being introduced into the plethysmograph. All odors were handled within independent tubing to prevent cross-contamination. Following each odor presentation, odor-vaporized air was passively cleared from the plethysmograph through an exhaust outlet at the chamber’s ceiling since the filtered room air is continuously delivered. Serving as a non-olfactory stimulus, a vibration motor (3V, 12000 RPM; BestTong), referred to as “buzz,” was secured to the outside of the plethysmograph. During trials of pseudorandom stimulus presentations, each stimulus was presented for 2s with a 60s inter-trial interval.

### Fiber photometric recording in head-fixed mice

#### Virus injection and optical fiber implant

Mice for this method were also used for data collection and their results published (albeit from different stimulus conditions wherein analyses were performed on well-learned odor-associated pairs) in a prior study^40^. As we previously described in that study, *D1-Cre* and *D2-Cre* mice were injected in the TuS with 1µL AAV5-EF1α-DIO-GCaMP6f-WPRE (obtained from the Penn Vector Core in the Gene Therapy Program of the University of Pennsylvania, 5.43×10^13^GC/mL). Viral injections were performed as described above (see ‘Animals’). Two weeks later, a 400µm core, 0.48NA optical fiber secured in a 2.5mm metal ferrule was chronically implanted to terminate in the TuS. A plastic head-bar was also affixed to the skull with dental cement for later head fixation. Animals were given a week for surgical recovery before undergoing behavioral testing.

#### In vivo fiber photometry

465nm (GCaMP excitation wavelength, driven at 210Hz) and 405nm (UV excitation wavelength, driven at 330Hz; control) light emitting diodes were coupled to a 5 port minicube (Doric lenses, FCM5) using 400µm, 0.48NA FCM optic fiber patch cords. Both excitation and emission light were directed through a patch cord connected to the animal by the implanted 2.5mm metal ferrule. Emission light was directed through the 5-port mini cube and coupled to femtowatt photoreceivers to monitor GFP and UV signals. The data were then digitized and pulse demodulated at 1kHz using a Tucker Davis Technologies processor (TDT). All parameters for imaging and acquisition of photometry data were consistent across D1- and D2-Cre mice.

#### Behavioral sessions

Animals were head-fixed and presented with sets of 3 novel odors, 4 times each. To enhance each animal’s contribution to a more rigorous data set, animals were tested a second time at least 24hrs later with sets of 3 different novel odors. Odors were delivered for 2s duration with a 17 ± 2s inter-trial interval through an odor port positioned 1cm from the animal’s snout.

#### Stimulus delivery

Odors were presented through an olfactometer and delivered through an odor port positioned 1cm from the animal’s snout. Odors included ethyl butyrate, 1,7-octadiene, isopentyl acetate, heptanal, 2-heptanone, R(+)-limonene, ethyl propionate, S(-)-limonene, methanol, methyl valerate, 2-butanone, 1,4-cineole, butanal, propyl acetate, allylbenzene, allyl bromide, isobutyl propionate, and 2-methylbutyraldehyde (Sigma Aldrich, all > 97% purity). Odors were diluted to 1Torr in mineral oil and further diluted by mixing 100mL odor vaporized N_2_ with 900mL medical grade N_2_ (Airgas). Stimuli were therefore presented at a total flow rate of 1L/min. Odor was continuously flowing to the odor port but removed by a vacuum prior to reaching the animal. Odor sets used varied across animals and across sessions.

#### Respiratory data acquisition

Respiration was recorded via an extranasal flow sensor (CPXL04GF, Honeywell) positioned near the animal’s nose as described^68^. A 1/8” inner diameter hose which terminated into the flow sensor was placed ∼7-10mm from the nose of the mouse during head fixation.

### Optogenetic stimulation in the plethysmograph

#### Virus injection and fiber implant

*DAT^IRES^-Cre x Ai9* mice or *VGluT2 fl/fl x TH-Cre* mice were injected unilaterally in the VTA with 400nL AAV5-EF1α-DIO-hChR2(E123T/T159C)-EYFP (Addgene 35509, titer: 1.3×10^13^vg/mL) or AAV5-EF1α-DIO-EYFP (for controls, UNC vector Core, titer: 3.5×10^12^vg/mL). While we used *DAT* to drive Cre expression in DAergic neurons in other experiments in this study, here we used the gene encoding tyrosine hydroxylase (*TH*) and specially *VGluT2 fl/fl x TH-Cre* mice since this line was already available in-house. While this *TH*-based strategy may not had captured the identical cell population as in the DAT-based strategy, and *VGluT2* may have been removed from non-DA cells in the VTA^114^, this mouse line allowed us to isolate the influence of DA on sniffing from glutamate. Viral injections were performed as previously described. In the same surgery, a 300µm core 0.39 NA optical fiber in a ceramic ferrule was chronically implanted to terminate in the TuS, NAcSh, or NAcC in *DAT^IRES^-Cre x Ai9* mice or in the TuS for *VGluT2 fl/fl x TH-Cre* and affixed to the skull with dental cement. Animals were tested no sooner than 4 weeks following surgery.

To reconstruct ChR2.EYFP injection sites in the VTA of *DAT^IRES^-Cre* mice (**Supplementary Fig 3**), the VTA was imaged from −2.92 to −3.80mm posterior bregma, as identified with use of a brain atlas^112^. An independent rater used Adobe Illutrator to trace density of expression across images.

#### Light stimulation and behavioral monitoring

Optogenetic stimulation (25Hz, 15ms pulse width, 1s duration, 18mW^3^) was delivered by a 473nm laser (CrystaLaser, DL473-100) and through a 300µm, 0.39NA fiber connected the laser to a fiber optic rotary joint, with a 400µm, 0.47NA fiber connecting the rotary joint to the mouse (brass mating sleeve). This paradigm was based on prior studies^39,40,54^ and pilot work done in the lab. Animals (*DAT^IRES^-Cre*: TuS *n*=11, NAcSh *n*=6, NAcC *n*=6, EYFP controls *n*=9; *VGluT2 fl/fl x TH-Cre*: TuS *n*=4, EYFP controls *n*=4) received one habituation day wherein they were gently tethered to the path cable and placed in the plethysmograph for 20mins. The following day, animals were again tethered, placed in the plethysmograph, and given a brief, 5min habituation period. Following this, photostimulations (7-10/mouse) were delivered with a 2 ± 1min inter-trial interval.

A subset of *DAT^IRES^-Cre* x *Ai9* (TuS: *n*=5, NAcSh: *n*=4, NAcC: *n*=4, EYFP controls: *n*=6) underwent testing with varying durations of photostimulation. Stimulations at 25Hz with a 15ms pulse width were delivered for durations of 120ms, 1s, 2s, or 4s. After a habituation day, mice received a single stimulation parameter per day over four consecutive days. Each mouse underwent the 1s duration on day 1, and the following 3 stimulation paradigms were randomized across mice over the remaining 3 testing days. Photostimulations (9-11/mouse) were delivered with a 2 ± 1min inter-trial interval.

Concluding testing, the same mice were then deeply anesthetized with urethane (1g/kg i.p.; Sigma Life Science), tethered, and placed in the plethysmograph and photostimulated as described above. Mice were transcardially perfused following recordings and within 30mins of urethane administration.

### Optogenetic inhibition in the plethysmograph

#### Virus injection and fiber implant

*DAT^IRES^-Cre x Ai9* mice or *DAT^IRES^-Cre* mice were injected bilaterally in the VTA with 300nL AAV5-EF1α-DIO-eNpHR 3.0-EYFP (Addgene 26966, titer: 1.0×10^13^ GC/mL). Viral injections were performed as previously described earlier in this study. In the same surgery, 400µm core, 0.39 NA optical fibers in ceramic ferrules were chronically implanted to bilaterally terminate in the TuS or NAcSh and affixed to the skull with dental cement. To afford precision in the bilateral spacing, the ferrules were first affixed in a 3D-printed adapter to ensure perfect 2.0mm from epicenter-to-epicenter spacing (tip-to-tip). Animals were tested no sooner than 4 weeks following surgery. Representative images of the primary injection site location of eNpHR 3.0-EYFP in the VTA were confirmed and imaged by an independent experimenter (**Supplementary Fig 7**).

#### Light stimulation and behavioral monitoring

Optogenetic stimulation (7-9mW^3^, 5-7s duration, constant, see Results for rationale) was delivered by 405nm or 560nm LEDs (ThorLabs: Lx405 and Lx560). A 400µm, 0.48NA fiber connected the LED to the mouse.

To examine whether optogenetic inhibition of DAergic terminals in the ventral striatum had any effect on odor- or buzz-evoked sniffing, animals then underwent a similar behavioral session as previously described (see Fiber photometric recording in the plethysmograph, Behavioral monitoring). Animals first received one habituation day wherein they were gently tethered to the path cable and placed in the plethysmograph for 20mins. Following this, in a counterbalanced, within-subjects design, animals were tested over a 2-week period wherein they received concurrent 405nm or 560nm light delivery (7s duration, constant), initiated at the onset of both odor and buzz delivery.

Separately, the same mice were again tested in a counterbalanced, within subject’s design over 2 days. Behavioral sessions lasted ∼10 minutes. Mice were gently tethered and placed in the plethysmograph. Starting at minute 2, 5s (constant) of 405nm or 560nm light was delivered every 20s for 8 minutes, for a total of 24 light deliveries.

### Machine learning to classify respiration

The study utilized a dataset comprising differently evoked sniffing data which consist of 261 samples, each 1s long. The dataset consisted of time series data representing respiratory frequencies (10ms bins).

Dynamic Time Warping (DTW) was employed as the distance metric for measuring the similarity between the time series data (respiratory frequency). DTW is known for its ability to align and compare time series data. DTW involves finding the optimal alignment between sets of data points of two sequences by warping the time axes. This allows the comparison of sequences with different lengths or different temporal structures. DTW creates a cost matrix, where each element represents the distance or dissimilarity between corresponding points in each of the two time series. Dynamic programming is used to efficiently find the optimal alignment by considering all possible alignments and selecting the one with the lowest cumulative distance ^59^

The k-Nearest Neighbors (k-NN) algorithm was utilized for classification of unknown data points based on the majority class of its k-nearest neighbors in the feature space. In the context of time series data, DTW can be used as a distance metric in the k-NN algorithm instead of using traditional Euclidean distance. For a given query time series, the algorithm identifies the k-NN from the training set based on DTW distance^60^. For each query time series, the k-NN were identified and a majority voting scheme was employed to assign the class label.

Prior to applying the DTW k-NN algorithm, time series data were applied min-max normalization to ensure a uniform scale across all features. Subsequently, the dataset was divided into training and testing sets, with 80% of the samples allocated for training (208 samples) and 20% for testing (53 samples). The performance of the DTW k-NN algorithm was evaluated in terms of its accuracy, precision, recall, and F1-score. The max warping window was set at 7 and k at 3 which identified the combination that maximized the performance metrics during experiments.

The experiments were conducted using Google Colab (Google Brain Team, Mountain View, CA, USA). The DTW k-NN algorithm^61^ was implemented in Python, utilizing the code available in the GitHub repository: https://github.com/markdregan/K-Nearest-Neighbors-with-Dynamic-Time-Warping. The dataset consisted of 63 photostimulation-evoked, 65 odor-evoked, 63 buzz-evoked, and 70 spontaneous respiratory/sniff bouts (*N*=261 total samples). SMOTE^62^ was applied to generate synthetic data samples to bootstrap all groups to a total of 70 samples.

### Single unit recordings of ventral striatum neuron activity in head-fixed mice

These mice were used for data collection and their results published in a prior study from our group, with exception of any neural analyses aligned to intranasal respiration as done herein^64^. As we described in that study, c57bl/6j mice were implanted with eight-channel tungsten electrode arrays in the TuS and implanted with intranasal cannula for simultaneous respiratory monitoring. A plastic head-bar was also affixed to the skull with dental cement for later head-fixation. Animals were given a week for surgical recovery before undergoing behavioral testing wherein water-motivated mice were shaped to lick for a fluid reward for one odor, not a different one. Each animal had 2-3 recording sessions on different days. Single neurons were sorted in Spike2, and then spiking and respiration were analyzed in MATLAB.

Units from the same animal but not from the same recording day were limited to different channels. The spontaneous period was within the intertrial interval excluding the five seconds from odor onset to avoid any swallowing or licking effect. The odor presentation period was defined as the time from odor onset to two seconds after odor onset. The respiration cycle was the interval between two positive peaks in respiration signals.

#### Sniffing burst detection

A high-pass filter (5 Hz) was applied to the respiration signal. The root mean square (RMS) was calculated using a 1s time window to quantify respiratory activity. The RMS was then transferred into the z-score. A threshold was set to 1 for sniffing onset detection.

#### Spike ratio and initial and terminal spike analyses

Respiration cycles were divided into four groups based on their duration, each containing 25% of the respiratory cycles. The spike number was then counted in each group for each unit. To establish control, 100 sets of randomly generated spikes were created, with the number of spikes in each set matching the actual spike number of the unit. The spike ratio was calculated as the spike number divided by the mean of the randomly generated spike number in each percentile group. The percentiles of the spontaneous and odor presentation periods were calculated separately. The initial spike was defined as the first spike without any spikes in the previous five respiration cycles. The terminal spike was the spike that had at least five spikes in the previous five respiration cycles and was not followed by any spikes in the five subsequent respiration cycles.

### Pharmacological manipulations

#### Intracranial cannulae implantation

Bilateral craniotomies were made to target the the TuS and NAcSh. Stainless steel bilateral guide cannulae (26GA, 2.0mm center-to-center distance, 5.3mm long from the base of the pedestal, P1 Technologies, Roanoke, VA) were lowered either to target the TuS (*n*=13) or the NAcSh (*n*=13). Cannulae were affixed to the skull via dental cement, and dummy cannulae with a 0.5mm projection from the end of the guide cannulae were then inserted and dust caps were added.

#### Selective antagonists and intracranial infusions

SCH23390 (Tocris Bioscience, Bristol, UK) was dissolved in 0.9% sterile saline to a concentration of 0.5µg/µL. 0.1µL was infused to deliver 0.05µg/hemisphere. Raclopride (Tocris Bioscience, Bristol, UK) was dissolved in 0.9% saline + 5% dimethyl sulfoxide to a concentration of 23.8µg/µL, and 0.21µL was infused to deliver 5µg/hemisphere. PG01037 (Tocris Bioscience, Bristol, UK) was dissolved in 0.9% saline + 12% DMSO to a concentration of 20µg/µL. 0.15µL was infused to achieve 3µg/hemisphere. Internal cannulae that were cut at lengths projecting 0.5mm past the guide cannulae and thus terminating in the TuS or NAc were connected to 25GA polyurethane tubing (Instech Laboratories, Plymouth Meeting, PA) backfilled with dH2O. 1µL Hamilton syringes (Reno, NV) were used to inject total volumes ranging from 0.1-0.21µL. Mice were gently scruffed, dust caps and dummy cannulae were removed, and the internal cannulae was inserted before mice were placed back in their home cages for infusions. All drugs were infused at a rate of 0.1µL/min followed by a 2min rest period before the internal cannulae was removed, dummy cannulae and dustcap replaced. For all groups, behavioral testing began 10min following the start of drug infusion. All mice were tested across a 6 week period and recieved all selective antagonists and their corresponding vehicles, with 7 days between testing sessions.

#### Behavioral methods

To monitor spontaneous and sensory-evoked sniffing behavior, mice were placed in the plethysmograph and underwent behavioral testing and stimulus delivery as previously described to monitor odor-, buzz-, and spontaneously-evoked sniffing (see Fiber photometric recording in the plethysmograph, Behavioral monitoring). Baseline breathing rates were identified as periods far removed from stimulation windows (*i.e.,* 2-5min, 14-17min, 30-33min).

### Histology

Mice receiving synaptophysin injections were perfused 3 weeks following injection. Mice undergoing behavior were perfused following the last day of behavioral monitoring. All mice were overdosed with sodium pentobarbital (Fatal-Plus, Patterson Veterinary) and transcardially perfused with 0.9% saline followed by 10% phosphate-buffered formalin. Brains were removed and stored (at 4°C) in a 10% formalin in 30% sucrose solution prior to sectioning. Brains were coronally sectioned at 40µm and sections were stored in Tris-buffered saline with 0.03% sodium azide. Microscopic examination of tissue occurred on a Ti2e inverted fluorescent microscope, and only mice with on-target manipulations (cannulae locations, viral injections, viral injections + optic fiber placements) contributed to data sets. Verification of brain regions and target sites was aided by use of a brain atlas^112^.

### Quantification and Statistical Analysis

#### Analysis of respiratory data

Inhalation peaks (sampled at 300Hz) were detected offline in Spike2 (Cambridge Electronic Design, Cambridge, UK). First, the respiratory data were filtered (0.1-12Hz band pass). Second, the maximum point of each respiratory cycle was identified, and instantaneous frequency was then calculated based off of the duration between one peak and its preceding peak. Data were down-sampled to 100Hz for subsequent analyses.

#### Analysis of fiber photometry data

Offline using a custom script in Spike 2, the GFP and UV signals were filtered (2^nd^ order, 25Hz low pass) and smoothed with a sliding window (9 bins averaged). A first-degree polynomial fit was used to find the relationship between the signal and control channels and generate a scaled control channel. The fitted control channel was subtracted from the signal channel and z-scored ΔF/F traces were calculated using a 3s pre-event baseline window (GRAB_DA_ analysis) or 2s pre-event baseline window (GCaMP6f for D1- and D2-Cre analysis).

Based on a series of pilot experiments, we found that on average, tethered animals in the plethysmograph typically do not perceive odors until 3s following odor control valve opening (as indicated by their reflexive/orienting sniffing responses). To account for this delay, the 3s after valve opening and odor onset serve as the pre-event baseline. Buzz-evoked GRAB_DA_ signals were analyzed relative to the buzz onset, and spontaneous sniffing was analyzed relative to bout onset.

Peak GRAB_DA_ signals and average sniff frequency were calculated within the 4s following odor analysis onset, 2s buzz duration, or spontaneous bout. Spontaneous sniffing bout durations varied and were identified based on prolonged periods of sniffing at frequencies ≥6Hz for ≥1s. Given the highly dynamic nature of sniffing and the rapid shift between respiratory frequencies in mice, we identified individual spontaneous sniff bouts wherein the mouse was sniffing at frequencies <6Hz for at least 2s prior to bout onset.

For D1- and D2-Cre GCaMP6f photometry, because animals were head-fixed and odor was delivered directly to the nose with high temporal precision, the 2s before odor delivery was used as the trial specific baseline for the z-scored ΔF/F values.

#### Additional statistical methods

Data were analyzed in GraphPad Prism. All performed statistical tests were two-sided. Sample sizes are consistent with those reported in the field. All data are reported as mean ± SEM unless otherwise reported. Specific tests and *p* values used can be found in the Results section or the figure legends.

## Acknowledgements

We thank Jacqueline Sipio for help with histology and Ellyse Thomas for expert animal care assistance.

This work was supported by R01DC014443 and R01DC016519 to D.W.W., R01DA049545, R01DA049449 and R01NS117061 to D.W.W. and M.M., R01DC006213 to M.M., and R00HL159232 to A.G.V.. N.L.J. was supported by NIDCD T32015994 and F31DC020364.

## Author Contributions

Conceptualization: NLJ & DWW

Methodology: NLJ, ACL, LAS, AGV, UMA, ZZ, MAG

Investigation: NLJ, ACL, LAS, ZZ, UMA, MAG

Visualization: NLJ, DWW, ACL

Funding acquisition: NLJ, DWW, MM, AGV

Supervision: DWW

Writing – original draft: NLJ

Writing – review & editing: NLJ, DWW, AGV, MM, UMA, LAS, ACL, ZZ

---

**Supp Figure 1:**
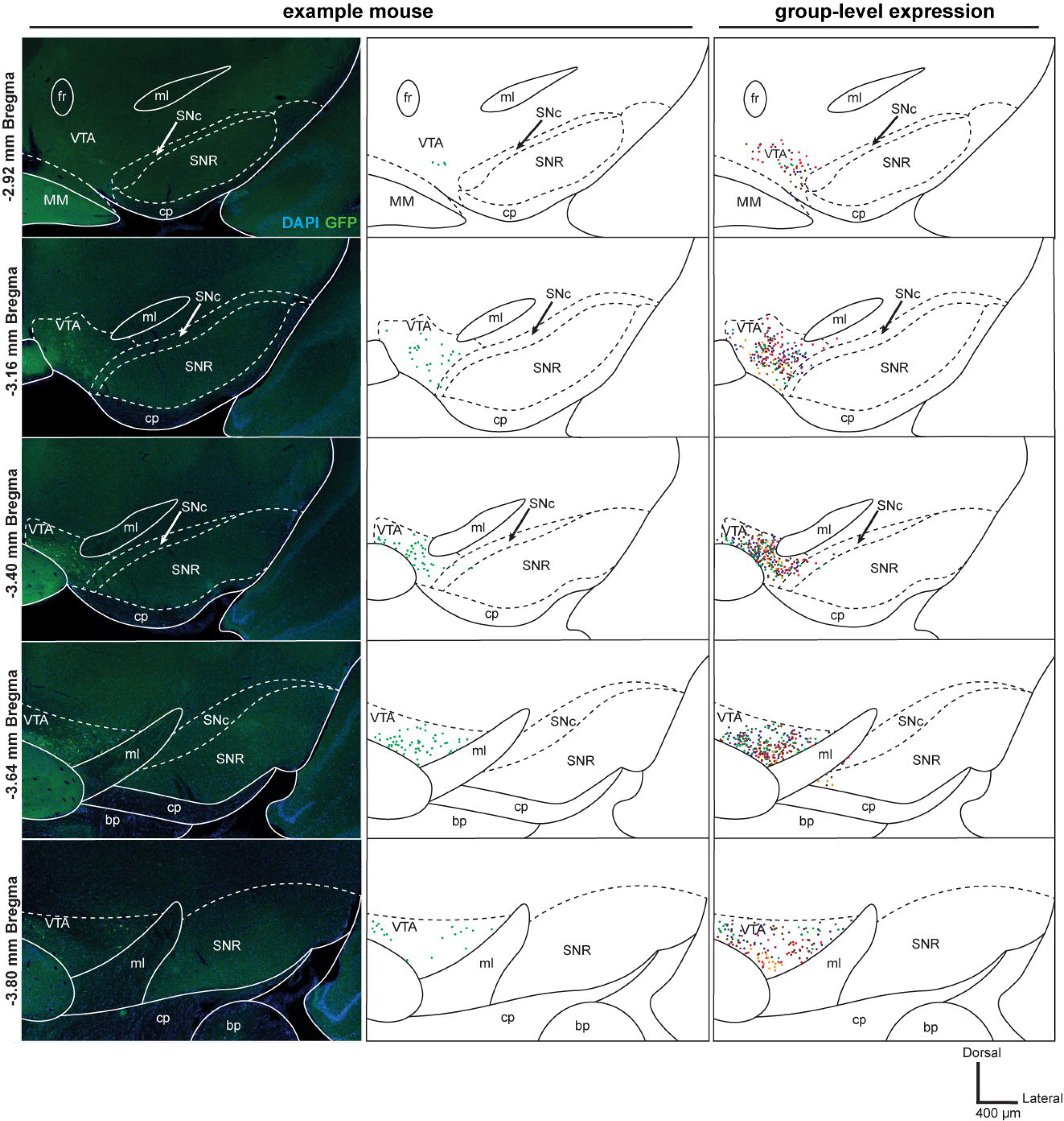

**Supp Figure 2:**
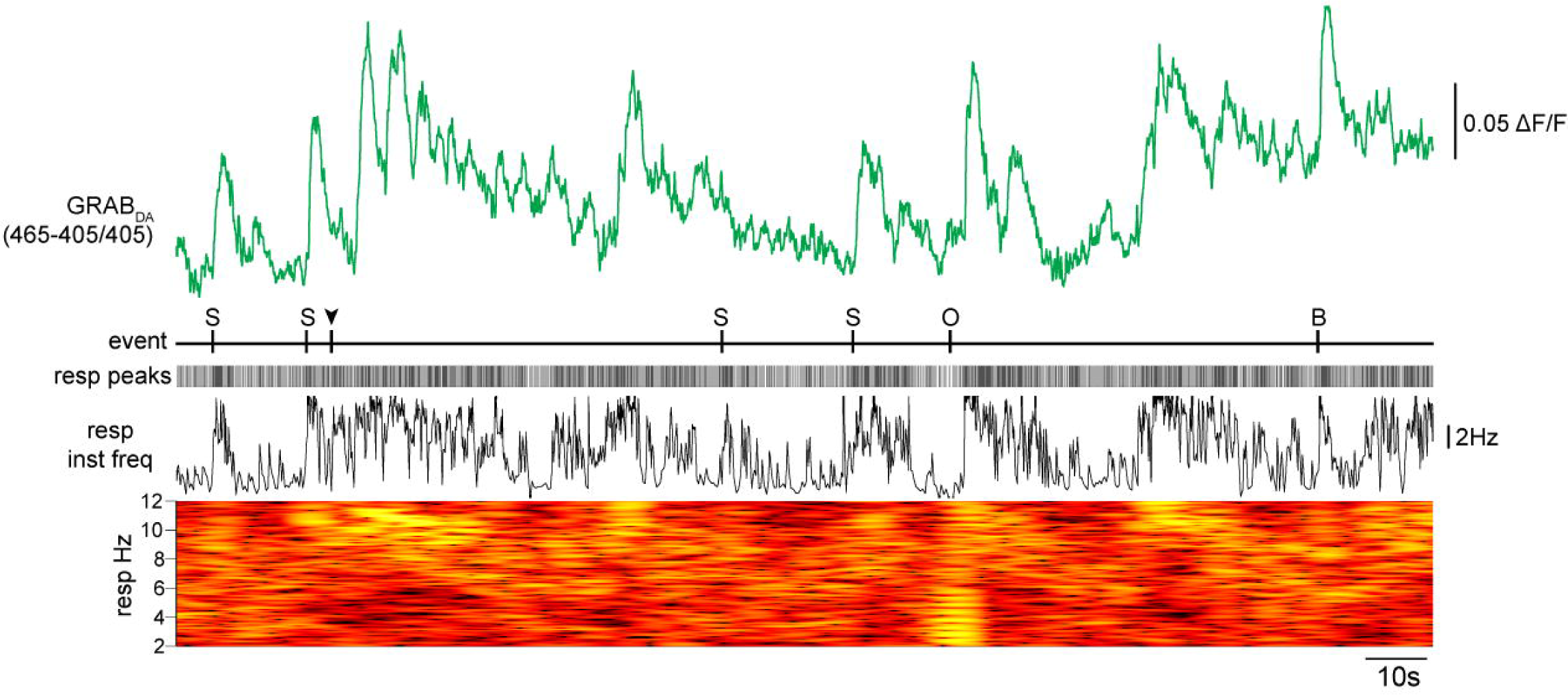

**Supp Figure 3:**
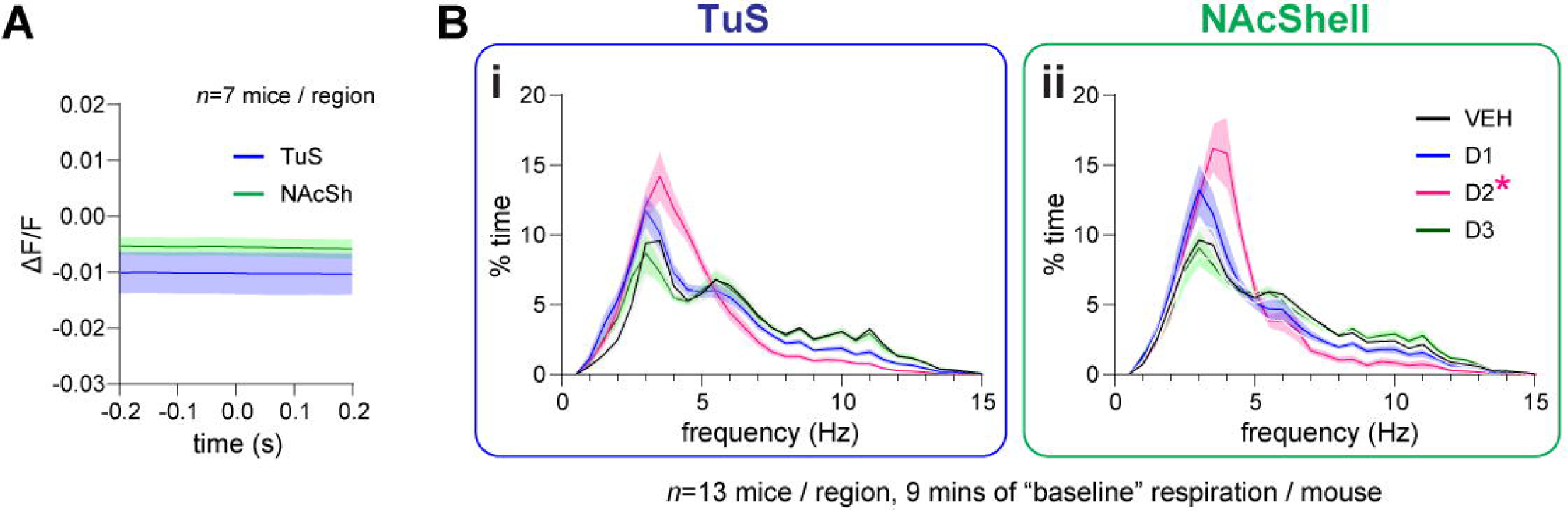

**Supp Figure 4:**
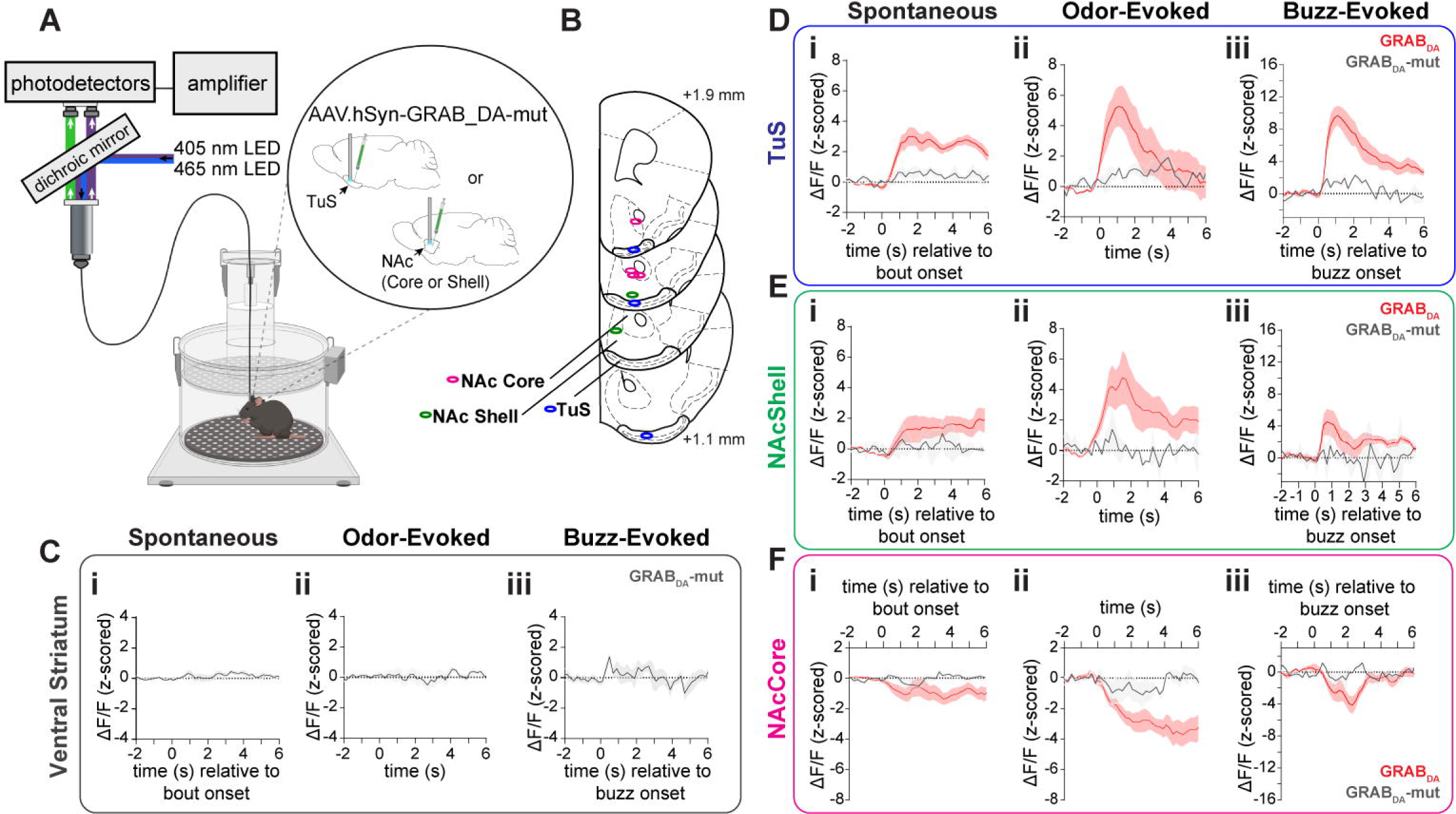

**Supp Figure 5:**
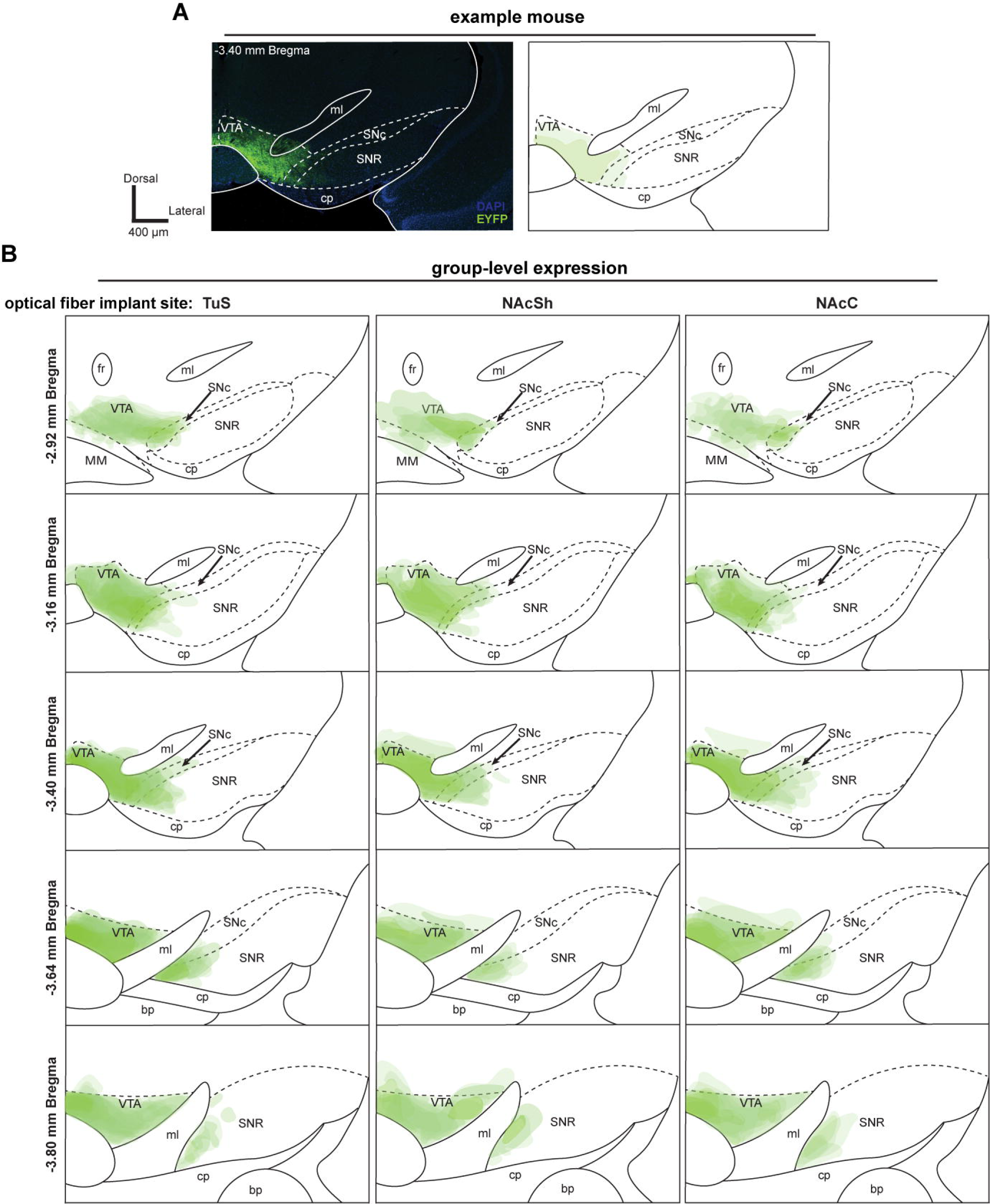

**Supp Figure 6:**
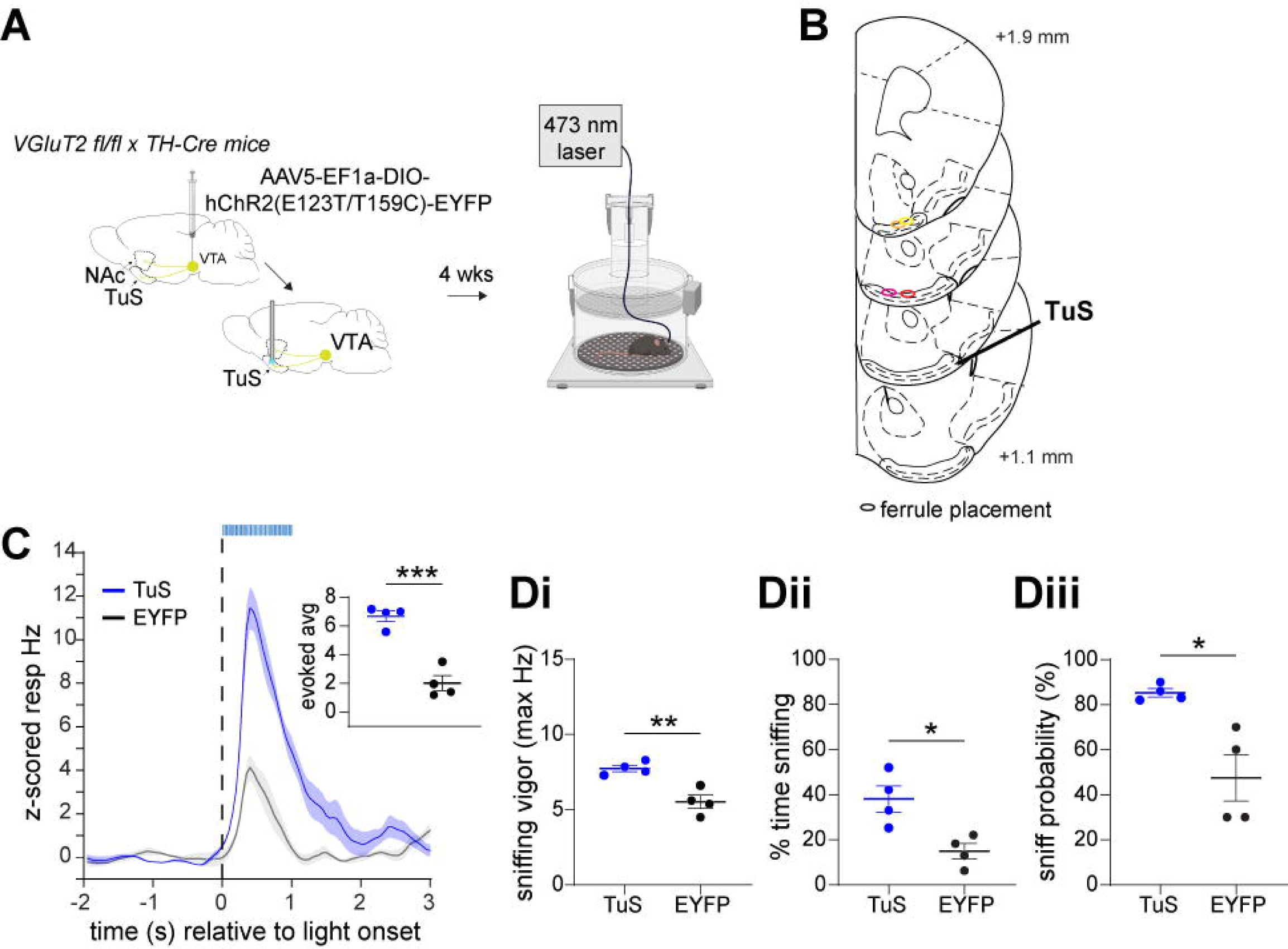

**Supp Figure 7:**
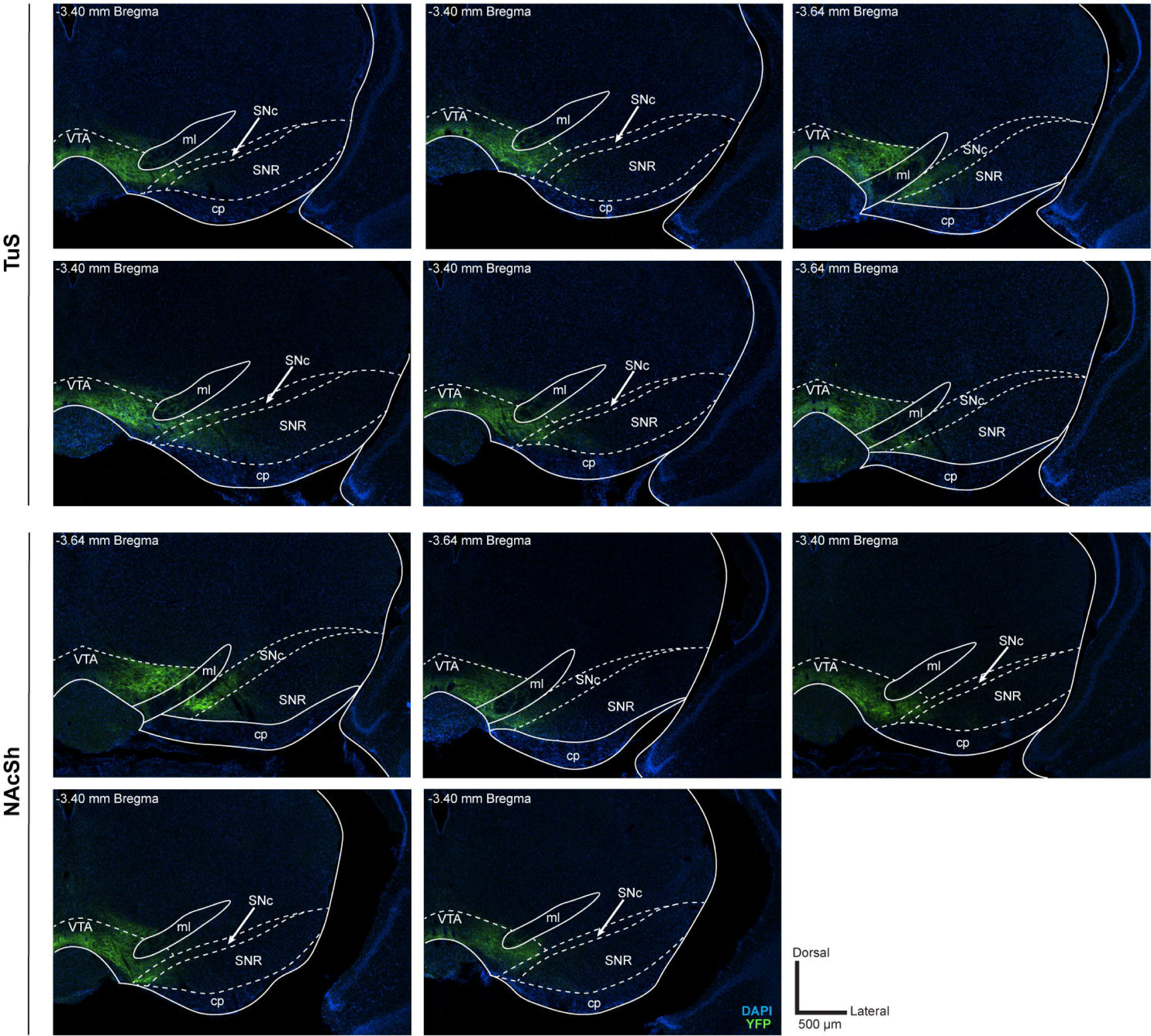

**Supp Figure 8:**
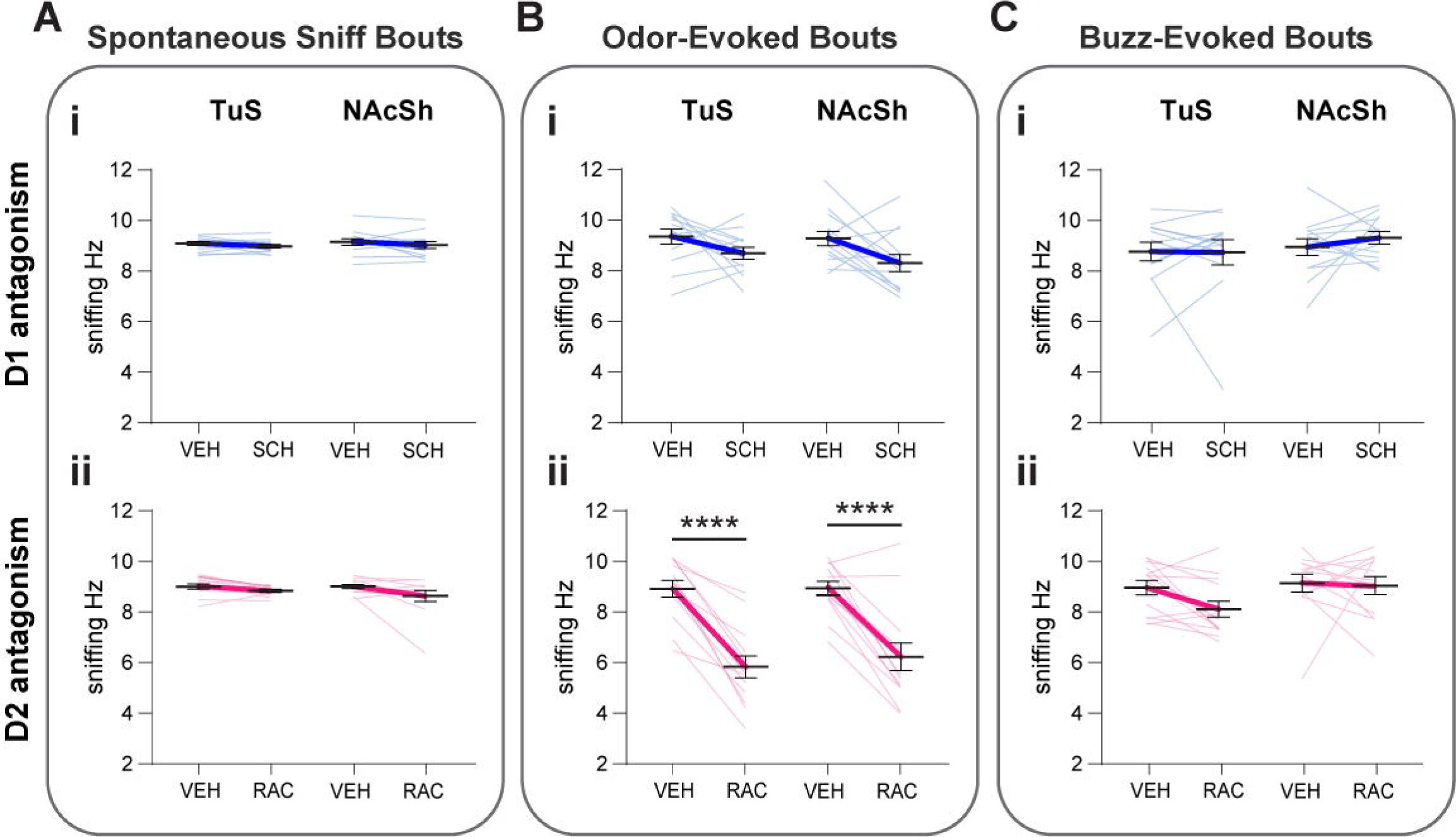

