## Supplemental Figure legends for "Sniffing can be initiated by dopamine’s actions on ventral striatum neurons"

**Supplementary Figure 1. Mapping of synaptophysin.mRuby expression in midbrain. Related to Figure 1.** Schematic depicting synaptophysin.mRuby injection sites across the anterior to posterior span of the VTA (-2.92 to -3.80mm posterior Bregma). *$DAT^{IRES}$-Cre* mice were injected in the VTA with AAV.FLEX.hSyn.mGFP.synaptophysin.mRuby allowing for GFP labeling of DAT+ cell bodies. Columns 1-2 shows representative images from one mouse (left) and an example of the identified GFP labeling in a corresponding atlas schematic (right). Column 3 shows a compilation of GFP labeled cell bodies across all animals (*n*=6). Groups of colored dots represent individual animal data. Abbreviations: VTA (ventral tegmental area), MM (medial mammillary nucleus), SNc (substantia nigra pars compacta), SNR (substantia nigra pars reticulata), fr (fasciculus retroflexus), ml (medial lemniscus), cp (cerebral peduncle), bp (brachium pontis).

**Supplementary Figure 2. Example DA dynamics during spontaneous and stimulus-evoked sniffing. Related to Figures 2 & 3.** Top shows a representative ΔF/F trace (200s; 465-405/405) during user-identified spontaneous sniffing bouts (S), experimentally delivered odors (O), and experimentally delivered multisensory buzz stimulus (B) from an animal with $GRAB_{DA}$ being monitored from within the TuS. Major rises in the $GRAB_{DA}$ signal are notable in periods coinciding with elevations in sniffing behavior, although in some instances there are sniff bouts that do not drive an appreciable rise in DA levels (downward arrowhead). Respiratory peaks (resp peaks) depict the peak of inhalation cycles and subsequent interpolated frequency trace shown beneath in black. Bottom sonogram displays respiratory power calculated by fast Fourier transform (FFT, with Hanning window, greatest power = yellow (70dB)).

**Supplementary Figure 3. No temporal linkage between ventral striatum DA levels, and subtle influence of DA receptor antagonism, on resting respiration. Related to Figures 2, 3, & 9. A.** Average $GRAB_{DA}$ ΔF/F signal aligned to individual breaths during periods of respiration where the mouse is at rest (*viz.,* lower frequency respiration (2-4Hz) and during periods far removed from stimulus delivery). For this we subtracted a 53ms lag from the two signals, like we did for **Figures 2 and 3** at 28ms, but here using the longer lag to account for how slow respiration lags when measured with a plethysmograph versus directly from within the nose (via cannula). Neither the $GRAB_{DA}$ signal in the TuS or NAc displayed any change relative to respiratory peaks (time 0s) when animals were simply engaged in resting/basal respiration **B.** Histogram of respiratory frequencies during periods far removed from stimulus delivery (9 mins/mouse) in vehicle-treated vs. selective DA receptor antagonist-treated mice in the TuS (Bi) and NAcSh (Bii). Vehicle data collapsed across groups for display only. Antagonism of D2 receptors in the NAcSh resulted in more occurrences of low frequency respiration (Wilcoxon signed rank: NAcSh *W*=-279, *p*=0.012; TuS *W*=-171, *p*=0.130), yet there were no effects of antagonizing D1 or D3 ventral striatum DA receptors (Wilcoxon signed rank: NAcSh D1 *W*=-175, *p*=0.121; NAcSh D3 *W*=17, *p*=0.888; TuS D1 *W*=-161 *p*=0.155; TuS D3 *W*=107, *p*=0.348). For statistical analyses, groups compared to within subject vehicles.

**Supplementary Figure 4. No overt sniffing-associated changes in $GRAB_{DA}$-mut signals. Related to Figures 2 & 3. A**. Schematic of fiber photometry and plethysmograph system used to simultaneously record $GRAB_{DA}$-mut signals and sniffing behavior. Animals were injected with AAV.hSyn-GRAB_DA-mut and implanted with an optic fiber in the TuS, NAcSh, or NAcC. 3 weeks later, animals were placed in the plethysmograph and recordings took place. Image made with BioRender. **D.** Optic fiber implant locations in the TuS, NAcSh, and NAcC (*n*=2-4/group).

**C.** Averaged z-scored ΔF/F traces of $GRAB_{DA}$-mut across the entire ventral striatum (averaged across TuS, NacC, and NAcSh) during spontaneous sniffing (i), odor-evoked sniffing (ii), and buzz-evoked sniffing (iii). No significant difference in average $GRAB_{DA}$-mut levels between 3s baseline and spontaneous sniffing, nor between 3s baseline and trial #1 of odor or buzz-evoked sniffing (spontaneous paired $t(8)=0.68$, $p=0.515$; odor paired $t(8)=0.05$, $p=0.958$, buzz paired $t(8)=0.586$, $p=0.574$). **D-F.** Averaged z-scored ΔF/F traces of $GRAB_{DA}$-mut (gray) in the TuS (D), NAcSh (E), and NAcC (F) during spontaneous (i), trial #1 of odor-evoked (ii), and trial #1 of buzz-evoked (iii) sniffing bouts compared to $GRAB_{DA}$ (red) levels from **Figures 2 and 3.**

**Supplementary Figure 5. Mapping of ChR2.EYFP expression in the midbrain. Related to Figures 4 & 5. A.** Example showing how regions of ChR2.EYFP expression were identified. *DAT*$^{IRES}$*-Cre x Ai9* mice were injected with AAV5-EF1α-DIO-hChRs(E123T/T159C)-EYFP in the VTA. Representative image from one mouse (left) with corresponding atlas schematic of that section (right). Darker green represents areas of greater ChR2.EYFP expression, while the lighter green represents areas of less expression (see Methods). **B.** Schematic reconstruction of ChR2.EYFP expression across the anterior to posterior span of the VTA (-2.92 – 3.80mm posterior Bregma). Columns 1-3 shows injection illustrations from animals whose optic fibers terminated in the TuS (left, $n$=11), NAcC (middle, $n$=6) and NAcSh (right, $n$=6). Group data were created by overlaying individual data from each animal. Darker green coloring represents regions with greater overlap in expression across animals.

**Supplementary Figure 6. Preservation of evoked sniffing in *VGluT2 fl/fl x TH-Cre* mice. Related to Figures 4 & 5. A.** Schematic of optogenetically-evoked sniffing in mice lacking the vesicular glutamate transporter VGluT2 in TH+ neurons. *VGluT2 fl/fl x TH-Cre* mice were injected with AAV5-EF1α-DIO-hChR2(E123T/T159C)-EYFP in the VTA, and optic fibers terminating in the TuS were implanted in the same surgery ($n$=4). **B**. Optic fiber implant locations in the TuS. **C.** Averaged z-scored respiratory frequency ± SEM of photostimulation-evoked sniffing. Stimulation onset denoted by vertical dotted black line and 25Hz, 1s duration by horizontal blue line. Inset denotes the evoked z-score average during the 1s stimulation ***$p<0.001$; unpaired $t(6)=7.31$, $p=0.0003$. **D.** Parameters of optogenetically-evoked sniffing during 1s photostimulation in ChR2 mice or EYFP controls. All data presented as mean ± SEM, points = individual mice. **$p<0.01$, *$p<0.05$. **Di.** Mean sniffing frequency (max sniffing frequency reached during photostimulation); unpaired $t(6)=4.57$, $p=0.004$. **Dii.** Mean percentage of time spent sniffing; unpaired $t(6)=3.45$ $p=0.014$. **Diii.** Mean probability that photostimulation will evoke sniffing; unpaired $t(6)=3.61$ $p=0.011$.

**Supplementary Figure 7. Examples of eNpHR3.0 expression in the midbrain. Related to Figure 6.** Representative images of one injection site for all mice used in the optical inhibition experiment (**Fig. 6**). *DAT*$^{IRES}$*-Cre* or *DAT*$^{IRES}$*-Cre x Ai9* mice were injected with AAV-Ef1α-DIO-eNpHR3.0-EYFP bilaterally in the VTA. A representative image of one injection site for each mouse was taken and overlaid with a corresponding atlas schematic of that section. Rows 1 and 2 are representative images from animals whose optic fibers terminated in the TuS ($n$=6) while rows 3 and 4 are representative images from animals whose optic fibers terminated in the NAcSh ($n$=5).

**Supplementary Figure 8. DA receptor antagonism reduces the frequency of sniffing. Related to Figure 9. A.** Impact of selective antagonism of D1 (Ai) or D2 (Aii) receptors on spontaneous sniffing frequency (average sniff frequency during spontaneous bouts). **B.** Impact of selective antagonism of D1 (Bi) or D2 (Bii) on sniffing frequency during novel odor presentations. **C.** Impact of selective antagonism of D1 (Ci) or D2 (Cii) on sniffing frequency during novel buzz presentation.
